# Pigment and isoprostanoid responses to cold atmospheric plasma-induced oxidative conditions in a natural epipelic diatom assemblage

**DOI:** 10.64898/2026.09.13.751231

**Authors:** Alexandre Desparmet, Camille Oger, Claire Vigor, Thierry Durand, Guillaume Reversat, Valérie Gros, Thierry Dufour, Cédric Hubas

## Abstract

Intertidal mudflats hosting diatom-dominated microphytobenthic communities experience strong variability in environmental conditions, which can lead to cellular oxidative stress. These constraints can, in some cases, affect diatom metabolism and photosynthetic performance in the short term, potentially compromising their survival and even leading to cell death. This study investigates pigment, isoprostanoid, and photophysiological changes in a diatom assemblage originating from a natural microphytobenthic biofilm under different durations of helium-oxygen bubbling cold atmospheric plasma experimental treatments, mimicking complex oxidative conditions of the environment. To this end, 2-min and 5-min treatments were applied to the diatom assemblage, and metabolic parameters were monitored over time up to +180 min after plasma treatment cessation. Despite physicochemical changes and slight perturbations in metabolite profiles induced by plasma treatments, photosynthetic capacities of the diatoms remained largely preserved. The duration-dependent effects of plasma treatments induced selective and mostly reversible changes, primarily characterized by an increase in the relative proportions of pheopigments and a decrease in the relative proportions of β-carotenes following the 5-min treatment. A transient decrease in the relative proportions of the isoprostanoids 10-F_4t_-neuroP and 5-F_2t_-IsoP was also observed. By exposing natural diatom assemblage to plasma-generated oxidative conditions, this study provides novel insights into how marine microbiome respond to oxidative perturbations, contributing to our understanding of diatom success in fluctuating intertidal environments.

## Introduction

Intertidal mudflats host highly productive microphytobenthic communities, often dominated by motile epipelic diatoms that form a three-dimensional structured, mucilaginous biofilms on muddy sediments (Underwood GJC & Barnett, M, 2006; Hubas et al., 2018; Desparmet et al., 2026a). These communities experience strong spatiotemporal fluctuations in environmental conditions, including irradiance, temperature, pH, nutrient availability, and desiccation state, together with biotic interactions that can impose multiple and concurrent physiological constraints (Underwood et al., 2022). Such conditions can promote the transient endogenous production and accumulation of reactive oxygen species (ROS) within diatoms, as excess irradiance can exceed oxygenic photosynthetic capacity and overstimulate the electron transport chain, leading to the production of singlet oxygen (^1^O_2_), superoxide anion (O_2_^•-^), hydroxyl radicals (•OH), and hydrogen peroxide (H_2_O_2_), natural by-products of oxygen metabolism and markers of cellular oxidative status (Waring et al., 2010; Lepetit & Dietzel, 2015; Foyer & Hanke, 2022). ROS may also arise directly from exogenous environmental sources, including precipitation, photooxidation of dissolved organic matter, and release from neighboring cells (Abele-Oeschger et al., 1997; Mittler et al., 2011; Diaz & Plummer, 2018; Morris et al., 2022; Cheng et al., 2025).

Depending on their concentration, reactivity, and localization, these highly reactive species can induce cellular and behavioural signaling responses (Lepetit & Dietzel, 2015; Carrara et al., 2021; Flori et al., 2025; Desparmet et al., 2026a), enabling diatoms to acclimate, adapt, and survive, while also interacting with biomolecules, oxidizing cellular components, disrupting cellular homeostasis and photosynthetic performance, and ultimately leading to cell death (Nishiyama et al., 2006, 2011; Foyer & Hanke, 2022). Diatoms possess diverse mechanisms to prevent and regulate oxidative stress, including enzymatic and non-enzymatic antioxidant defenses, photoprotective xanthophyll-cycle pigments, and behavioural responses (Nymark et al., 2009; Waring et al., 2010; Lepetit & Dietzel, 2015; Serôdio et al., 2012; Desparmet et al., 2026a), which together contribute to their persistence in variable environments and partly explain their ecological success (Serôdio & Lavaud, 2020; Pierella Karlusich et al., 2025).

Among the cellular components potentially affected by ROS, polyunsaturated fatty acids are particularly susceptible to ROS-mediated peroxidation through hydrogen atom abstraction from bis-allylic positions, generating a diverse range of oxygenated lipid secondary metabolites, collectively referred to as oxylipins (Galano et al., 2017; Morris et al., 2022; Knieper et al., 2023). Widely reported across living organisms, including fungi, plants, animals, bacteria, cyanobacteria, and microalgae rich in precursors such as eicosapentaenoic acid in diatoms, oxygen incorporation into the carbon backbone of lipids to form oxylipins can occur either through regulated enzymatic processes or through non-enzymatic, uncontrolled oxidation (Galano et al., 2017; Beccaccioli et al., 2022; Doose et al., 2024; Desparmet et al., 2026b).

Enzymatically derived diatom oxylipins are increasingly recognized as biologically active mediators that can be interpreted by cells to modulate meaningful biological responses, including antibacterial and antipredator defenses, such as defense against copepod grazing (Miralto et al., 1999; Ianora et al., 2004; Ianora & Miralto, 2010; Orefice et al., 2022), modulation of defense-related gene expression (Sabharwal et al., 2017; Knieper et al., 2023), and allelochemical interactions involved in interspecific competition, as well as inter-and intracellular communication and signaling, including interactions with cellular calcium signaling pathways (Vardi et al., 2006; Nanjappa et al., 2014; Di Dato et al., 2020; Orefice et al., 2022). In contrast, non-enzymatic oxylipins, called isoprostanoids, arise from free-radical-mediated lipid peroxidation and were mainly considered passive biomarkers of oxidative damage, with their formation depending mainly on the availability and physicochemical properties of lipid precursors, their localization relative to ROS sources, and ROS concentrations (Jahn et al., 2008; Galano et al., 2017; Ahmed et al., 2020). Nevertheless, in recent years, isoprostanoids have undergone a shift in perspective and are now increasingly recognized as a family of bioactive metabolites, although their physiological roles remain less well characterized across living systems than those of enzymatically derived oxylipins (Bultel-Poncé et al., 2016; Galano et al., 2017). In microalgae, isoprostanoid profiles are more diverse but generally less abundant than enzymatically derived oxylipins (Linares-Maurizi et al., 2023), and their specific profiles have been proposed as potential chemotaxonomic tools for discriminating among diatom species (Lupette et al., 2018; Vigor et al., 2020; Linares-Maurizi et al., 2023, 2024).

Here, we investigated the effects of different durations of helium-oxygene bubbling cold atmospheric plasma (CAP) treatment on isoprostanoid and lipophilic pigment profiles, as well as photosynthetic performance, in a natural microphytobenthic diatom assemblage. More specifically, CAP is expected to generate short-(i.e., ^1^O_2_, O_2_^•-^, and •OH) and long-lived reactive species (i.e., H_2_O_2_): a complex exogenous oxidative environment utilized to compare diatoms’ responses with those previously observed under photooxidative stress. Since CAP does not impose any photosynthetic stress, we can therefore verify whether pigment and isoprostanoid profiles can reveal stress-related changes before substantial impairment of photosynthetic performance becomes detectable.

We hypothesized that increasing CAP exposure time would induce progressively stronger and transient alterations in pigment and isoprostanoid profiles, potentially accompanied by changes in photosynthetic performance. This experimental approach complements previous investigations of isoprostanoid responses to photooxidative stress in a comparable microphytobenthic diatom assemblage from the same sampling site (Doose et al., 2024).

## Materials and methods

### Experimental microphytobenthic community preparation

#### Biofilm sampling

Using a shovel, the upper centimeter of fine-grained sediments hosting a natural microphytobenthic community was collected at low tide from a disused outdoor breeding pond (Concarneau Marine Station, France; 47°52ʹ05.85ʺN, 3°55ʹ00.51ʺW; 18 June 2025, 5:30 PM). This pond is subject to a semi-diurnal tidal cycle (see Figs. 1, 2 Chapter 2) and retains at least 5 cm of clear water above the muddy surface at low tide, ensuring continuous immersion of the community. To allow biofilm reconstitution, samples were placed in plastic containers, transported to the laboratory, and allowed to settle in darkness (< 1 µmol photons m^-2^ s^-1^) at 15°C, matching *in situ* temperature.

**Figure 1.**
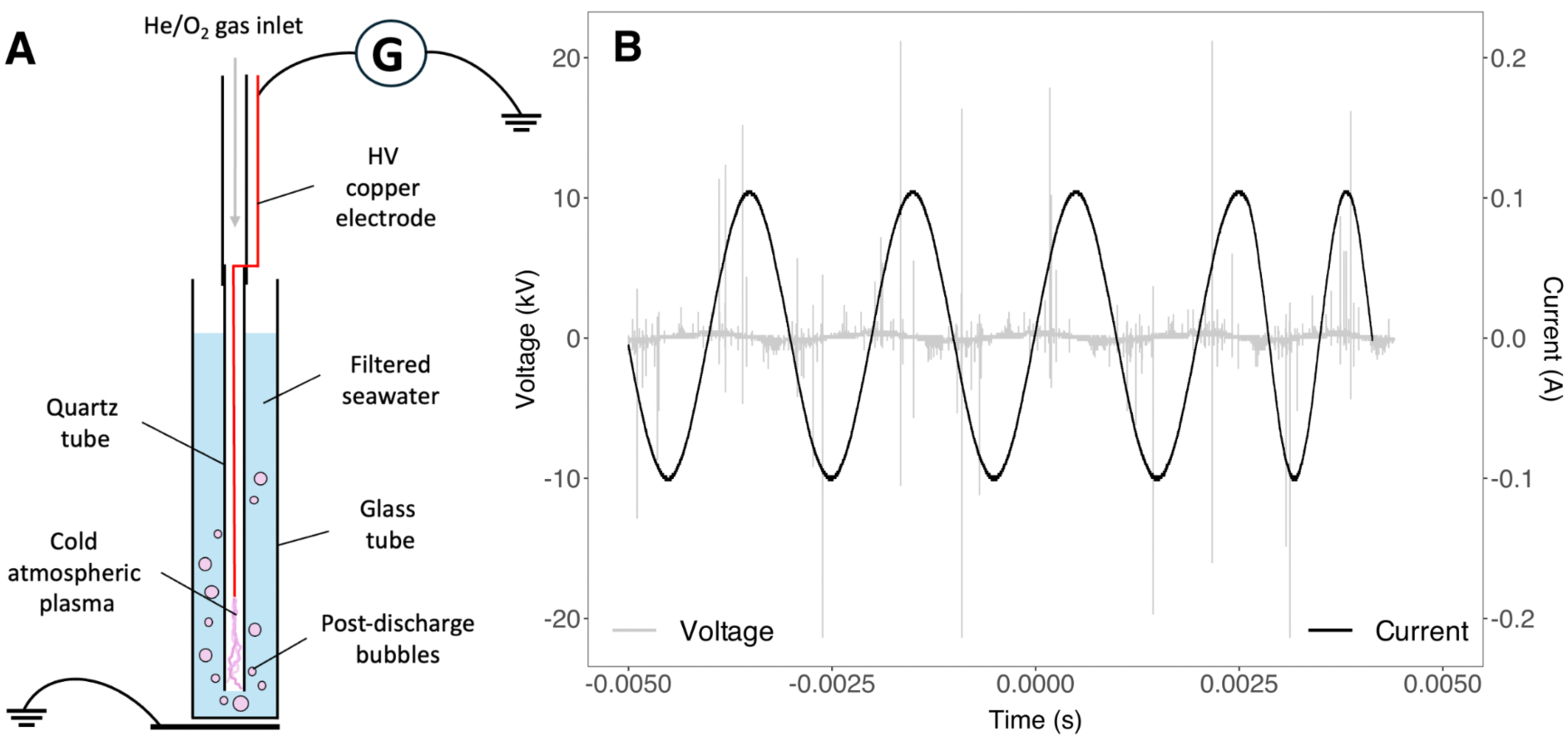
Experimental setup. **A.** Bubbling experimental cold atmospheric plasma setup used for microalgal suspension treatments; **B.** Cold atmospheric plasma electrical parameters applied to the copper electrode (A = 10 kV, 500 Hz).

#### Microalgal suspension and replicate conditioning

Following Desparmet et al. (2026a), a microalgal suspension was prepared the day prior to the experiment by transferring the upper millimeter of the reconstituted biofilm community into filtered seawater from the sampling site, using tweezers. Microalgal concentration was determined from five transects (250 grid cells; n _Total cell count_ = 733 diatoms) of a 1:4 diluted suspension using a 1 mL Sedgewick-Rafter counting chamber. Community composition was determined by identifying cells to the genus level by optical microscopy, and according to previous metabarcoding and scanning electron microscopy characterization (Desparmet et al., 2026a; Gaubert-Boussarie et al., 2020).

For each experimental treatment, replicates dedicated to pigment and isoprostanoid analyses (n = 4) consisted of 30 mL dark-adapted microalgal suspensions (< 1 µmol photons m^-2^ s^-1^) placed in glass tubes (77 mm height). Immediately after cold atmospheric plasma (CAP) treatment (see below), these metabolic replicates were frozen in liquid nitrogen, lyophilized, and stored at-80°C until extraction and analysis. Additional replicates (n = 3) were dedicated to photophysiological measurements.

### Experimental treatments

#### Cold atmospheric plasma generation and treatments

To generate a complex exogenous oxidative environment while avoiding direct light excitation, a helium-oxygen CAP discharge was applied to the microalgal suspension. CAP was generated inside a quartz tube reactor and coupled directly to the microalgal suspension by bubbling the plasma-activated gas through the submerged tube outlet (Fig. 1A). This bubbling configuration served two main purposes: (i) delivering reactive species directly into the microalgal suspension, thereby maximizing contact between the cells and short-lived species generated near the gas-liquid interface, and (ii) continuously resuspending the large benthic diatom cells to prevent their rapid sedimentation and ensure homogeneous CAP exposure.

The CAP was produced using a copper electrode connected to a high-voltage power supply (10 kV, 500 Hz; Fig. 1B), with a helium-oxygen (He/O_2_) carrier gas mixture flowing through the tube at controlled flow rates (He: 0.25 L min^-1^; O_2_: 10 cm^3^ min^-1^). The applied-voltage and discharge current waveforms were measured using a Teledyne LeCroy WaveSurfer 3054 digital oscilloscope, in combination with high-voltage probes (Tektronix P6015A 1000:1) and a Pearson 2877 current monitor.

Two CAP exposure durations, 2 min and 5 min, were tested as oxidative treatments. Negative controls (*i.e.* no CAP treatment and no gas bubbling) were performed in parallel to account for temporal metabolic changes unrelated to CAP treatment.

#### Characterization of plasma-generated conditions

The radiative components produced in the gaseous phase by the CAP were characterized by optical emission spectroscopy, using an Andor SR-750-B1-R spectrometer coupled to an Andor iStar intensified charge-coupled device camera Czerny-Turner configuration. The spectroradiometer had a focal length of 750 mm and was equipped with a 1200 groove mm^-1^ grating, blazed at 300 nm. For all experiments, parameters include an exposure time of 0.5 s, 10 accumulations and a gain level of 2000 a.u..

Changes in microalgal suspension temperature during CAP exposure were indirectly monitored in three independent experiments by measuring the apparent surface temperature of the reactor walls using a VarioCAM HD Head 680/30 thermographic camera (Jenoptik & InfraTec) with Ibris 3.1 Plus software, assuming rapid thermal equilibration between the reactor walls and the suspension. Indeed, although infrared thermography does not provide a direct measurement of the bulk suspension temperature, rapid heat exchange between the thin quartz wall and the surrounding continuously mixed suspension is expected to limit sustained temperature differences between the two compartments. Thus, as a first approximation, T_suspension_ can reasonably be assumed to approximate T_wall_, while acknowledging that local and transient temperature gradients cannot be excluded. Infrared images were acquired every 6 s from the beginning of exposure until 5 min. In parallel, the pH of the microalgal suspension was monitored (n = 3) using a Checker Plus HI98100 pH meter from Hanna Instruments.

#### Metabolic analysis and photosynthetic parameters

For both the 2-and 5-min CAP treatments, the photophysiological and metabolic, the temporal recovery of the microalgal metabolic response were assessed by monitoring at multiple time points: before CAP treatment, immediately after treatment cessation, and 30 and 180 minutes post-treatment (Table 1).

**Table 1.** Experimental setup. Summary of the materials and treatments tested in this study. Raw incident photosynthetic photon flux density (PPFD, in µmol photons m^-2^ s^-1^).

| Conditions | Dark-adapted control | Shorter CAP treatment | Longer CAP treatment |
| --- | --- | --- | --- |
| Description | Negative control<br>(0 min, dark-adapted<br>< 1 PPFD) | He/O<sub>2</sub> CAP<br>(2 min, dark-adapted<br>< 1 PPFD) | He/O<sub>2</sub> CAP<br>(5 min, dark-adapted<br>< 1 PPFD) |
| Experimental setup | - | He/O<sub>2</sub> CAP-activated seawater microalgal suspension through a copper electrode powered by a high-voltage power (10kV and 500 Hz) |  |
| Time-point monitoring | In parallel with treated replicates, at corresponding time points | Before CAP treatment, Immediately after CAP treatment, +30 min after CAP cessation, +180 min after CAP cessation |  |
| Metabolic analyses | Microalgal suspension, n = 4 |  |  |
| Photophysiological measurements | Microalgal suspension, n = 3 |  |  |

#### Lipophilic pigments analysis

Lipophilic pigments were extracted following the protocol of Brotas and Plante-Cuny (2003), then processed by high-performance liquid chromatography and quantified using calibration curves based on phytoplankton pigment standards from Desparmet et al. (2026a). For each replicate, 40 mg of freeze-dried material was sonicated for 30 s in 1 mL of 95% methanol buffered with 2% ammonium acetate, before incubation for 15 min in the dark at-21°C. The extract was then filtered through a 0.2 µm PTFE filter, and a 100 µL aliquot was injected into an Agilent 1260 Infinity high-performance liquid chromatography system equipped with a Supelcosil C18 reverse-phase column. The pigment separation method, identical to that used in Desparmet et al. (2026), employed a flow rate of 0.6 mL min^-1^ using a solvent mixture of A (0.5 M ammonium acetate in 85:15 methanol:water), B (90:10 acetonitrile:water), and C (100% ethyl acetate). Eluted lipophilic pigments were identified based on their retention times and absorption spectra, recorded with a UV-VIS photodiode array detector (DAD 1260 VL, 250 – 900 nm).

A total of 53 lipophilic pigments were detected and identified, and their relative proportions were calculated based on their respective concentrations. Two groups of derived chlorophyll *a* were identified: the “chlorophyll *a*-derivatives” group includes unidentified chlorophyll *a*-like pigments, as well as the allomer, and epimer forms; the “pheopigments” group includes pheophytins *a*, pheophorbides *a*, and pyropheophytin *a*. In addition, the diadinoxanthin de-epoxidation state (%), defined as the ratio of diatoxanthin to the diadinoxanthin + diatoxanthin pool, was calculated as [diatoxanthin]/([diadinoxanthin] + [diatoxanthin]) x100.

#### Isoprostanoid extraction and processing

Isoprostanoids were extracted following the method published by Vigor et al. (2018), as adapted by Doose et al. (2024). For each replicate, 200 mg of freeze-dried material was ground for 30 s in a lysing tube (Lysing Matrix D from MP Biomedical ref.6913500) at 6.5 m s^-1^ (FastPrep-24), in the presence of 25 μL of BHT (butylated hydroxytoluene, 1% in methanol), 1 mL of methanol, and 4 μL of internal standard mixture (MIX4IS; comprising C19, C21_15F_2t_IsoP, d4-10-epi-10-F_4t_-NeuroP, and d4_15F_2t_IsoP; 1 ng μL^-1^). The resulting homogenate was transferred into an empty tube, while the lysing tube was rinsed successively with 1 mL of methanol and 2 × 0.75 mL of phosphate buffer (pH 2, NaCl-saturated), with each rinse transferred sequentially into the same tube. This mixture was then agitated for 30 min using a GrantBio PTR35 multirotator (100 rpm 45 sec – 90° 15 sec) and then centrifuged at 4700 rpm using a FISHER GT2R Expert centrifuge equipped with a Thermo TX400 rotor (29 cm) for 5 min. Once recovered, the supernatant was combined with 4 mL of cold chloroform, vortexed for 30 s, and centrifuged at 2000 rpm for 5 min at 4°C using the same centrifuge. The lower organic phase was then collected and dried using a TurboVac apparatus at 50°C for 15–20 min (N_2_ flow rate: 0.5–1.2 mL min^-1^). Hydrolysis of the dried extracts was performed by adding 950 μL of 1 M KOH and incubating on a mixing plate at 40°C for 30 min, followed by the addition of 1 mL of 40 mM formic acid prior to solid-phase extraction on pre-conditioned Oasis mixed-mode polymeric sorbent cartridges (Oasis MAX Cartridge, 60 mg, Waters). Once samples were loaded onto the cartridges, undesired compounds were removed automatically using 1.5 mL of 2% (v/v) NH_3_, 1.5 mL of MeOH/20 mM formic acid (30:70; v/v), 1.5 mL of hexane, and 1.5 mL of hexane/ethyl acetate (70:30; v/v). Isoprostanoids were then eluted with 2 × 750 µL of a hexane/EtOH/acetic acid mixture (70:29.4:0.6; v/v/v). Samples were dried using a TurboVac at 50°C for 5 min (N_2_ flow rate: 0.5– 1.2 mL min^-1^), and the dried extracts were concentrated at the bottom of the tube by adding 200 µL of the hexane/EtOH/acetic acid mixture (70:29.4:0.6; v/v/v) and re-dried using a TurboVac at 50°C for 2 min (N_2_ flow rate: 0.5–1.2 mL min^-1^). Samples were reconstituted by adding 100 μL of mobile phase solvents (H₂O/ACN; 83:17; v/v), transferred into 0.45 μm filtered Eppendorf tubes (Nanosep Centrifugal Devices), and centrifuged at 10 000 rpm using a MICROSTAR 17R centrifuge (rotor diameter: 14 cm; ref.75003424 CH.50036PP) for 1 min at room temperature. The resulting aliquot was transferred into a HPLC analytical vial for further analysis. Quality control samples were also prepared by adding 4 μL of Prostamix GR57 SM0.5, containing all 54 oxylipin standards at 0.5 ng μL^-1^.

Compound separation was performed by injecting 5 μL of the extract into a micro-LC-MS/MS 5500 Q-Trap system from Sciex, combining high-performance liquid chromatography with tandem mass spectrometry. The mobile phases consisted of a gradient of H_2_O with 0.1% (v/v) HCO_2_H and ACN/MeOH (80:20, v/v), at a flow rate of 0.03 mL min^-1^, through a HALO C18 analytical column (100 × 0.5 mm, 2.7 μm; Eksigent Technologies, CA, USA) maintained at 40°C. The elution gradient was as follows: 17% B at 0 min, 17% B at 2.6 min, 21% B at 2.85 min, 25% B at 7.3 min, 28.5% B at 8.8 min, 33.3% B at 11 min, 40% B at 15 min, and 95% B from 16.5 to 18 min. Mass spectrometry analyses were performed in negative electrospray ionization (ESI) mode on an AB Sciex QTRAP 5500 (Sciex Applied Biosystems, ON, Canada). The source voltage was maintained at-4.5 kV, with nitrogen used as curtain gas (30 psi) and nebulizer gas (20 psi) at room temperature. To analyze the targeted compounds within a 90 s detection window, ionic fragmentation products of each deprotonated analyte [M–H]^-^ were monitored in multiple reaction monitoring (MRM) mode, using nitrogen as the collision gas. Two transitions per compound, one for quantification (T1) and one for identification (T2), were predetermined by MS/MS analysis of the corresponding standards. LC-MS/MS data acquisition was performed using Analyst® software (Sciex Applied Biosystems), and peak integration and analyte quantification were carried out using MultiQuant 3.0 software (Sciex Applied Biosystems).

#### Chlorophyll fluorescence: Pulse amplitude modulated fluorometry

Changes in photosynthetic parameters under treatments were assessed by measuring chlorophyll *a* fluorescence using the MonitoringPen MP 100-E (Photon System Instruments). For each replicate, an 8 mL dark-adapted microalgal suspension volume, taken from the 30 mL experimental volume, was transferred into a 6-well plate to undergo a rapid light curve protocol consisting of seven 1-min irradiance steps, increasing from 10 to 1000 µmol photons m^-2^ s^-1^ (λ _LED excitation_ = 470 nm), with consistent gain, superpulse, and flashpulse intensity settings maintained across all photophysiological measurements. The relative electron transport rate (rETR) of photosystem II (PSII), corresponding to values extracted directly from the light curve, reflects photosynthetic productivity (Perkins et al., 2010) and, in some cases, serves as a proxy for carbon fixation (Barranguet & Kromkamp, 2000). The initial linear α-slope, representing the maximum light-use efficiency of photosystem II, was derived from rETR-irradiance (rETR-E) curves using a previously described and modified model (Eilers & Peeters, 1988; Silsbe & Kromkamp, 2012). The maximum PSII quantum efficiency of dark-adapted cells (Fv/Fm), together with its regulated (Y(NPQ)) and non-regulated (Y(NO)) non-photochemical quenching components, was calculated following the equations of Consalvey et al. (2005) and Klughammer & Schreiber (2008) (Supplementary Table 1). Photosynthetic measurements (n = 3) were performed in parallel with metabolic replicates, at corresponding time points (Table 1).

#### Cell viability

The nuclear stain Sytox Green (ref. S7020) from ThermoFisher, which selectively penetrates cells with compromised membrane permeability, was used to assess cell viability, as commonly applied in studies on various microalgae (Veldhuis et al., 2001; Casotti et al., 2005; Vardi et al., 2006; Goessling et al., 2016; Volpert et al., 2018). For treated-samples, 200 µL of the CAP-treated microalgal suspension was immediately transferred into a 96-well plate, followed by the addition of 100 µL of filtered seawater from the sampling site and 35 µL of 50 µM SYTOX Green solution, prepared by diluting a 5 mM stock solution in the same filtered seawater. The seawater used was filtered through a sand filter and subsequently treated with ultraviolet light (pH_filtered seawater_ = 7.9). In parallel, positive and negative controls, as well as a blank consisting 300 µL of filtered seawater, were run. The negative control consisted of 200 µL of untreated microalgal suspension mixed with 100 µL of filtered seawater and 35 µL of 50 µM SYTOX Green, to confirm that the probe did not penetrate cells with uncompromised membrane integrity and stain their nuclear DNA. Conversely, the positive control consisted of 200 µL of untreated suspension combined with 100 µL of 70% EtOH and 35 µL of 50 µM SYTOX Green, to confirm the ability of the probe to penetrate cells with compromised membrane integrity and stain their nuclear DNA. The final SYTOX Green concentration in each well was approximately 5.2 µM. Fluorescence of stained nuclei was acquired at 4× magnification under GFP mode wavelengths (λ _LED excitation_ = 469 nm and λ _LED emission_ = 525 nm), after a 180 min incubation period, using a Cytation 3 cell imaging plate reader (BioTek).

#### Data treatment

Relative lipophilic pigment proportions were calculated based on their concentrations, whereas relative isoprostanoid proportions were calculated based on their raw peak areas obtained from chromatograms.

Pigment and isoprostanoid relative proportion data were analysed as two data blocks using multiple factor analysis (MFA) after Hellinger transformation. Euclidean distances between group centroids in the MFA space were calculated using the coordinates across all MFA dimensions and visualised as a heatmap to assess multivariate dissimilarities among experimental groups. Subsequently, a between-class multiple factor analysis (BC-MFA) was performed to assess the extent to which the multivariate inertia of the combined data blocks was structured by the predefined experimental groups. The significance of the between-group structure was assessed using a Monte-Carlo permutation test.

Two separate two-way permutational multivariate analysis of variance (PERMANOVA) were performed on the pigment and isoprostanoid datasets to assess the effects of time factor, experimental condition factor, and their interaction on multivariate variation in relative compound proportions, using Bray-Curtis dissimilarities. Homogeneity of multivariate dispersions among experimental groups was assessed using a permutation test to evaluate whether differences in dispersion could affect the interpretation of PERMANOVA results.

To assess differences among experimental groups, changes in individual compound and photophysiological parameters were tested using the Van der Waerden test.

## Results

### Cold atmospheric plasma physicochemical-induced changes

The experimental configuration allowed the plasma-activated He/O_2_ gas mixture to be bubbled directly through the microalgal suspension. This maintained continuous resuspension of the microalgae, thereby limiting diatom sedimentation and promoting contact between the cells and CAP-generated species transferred to the liquid phase. The OES spectrum contained several features attributable to electronically excited molecular and atomic species (Fig. 2 and Table 2).

**Figure 2.**
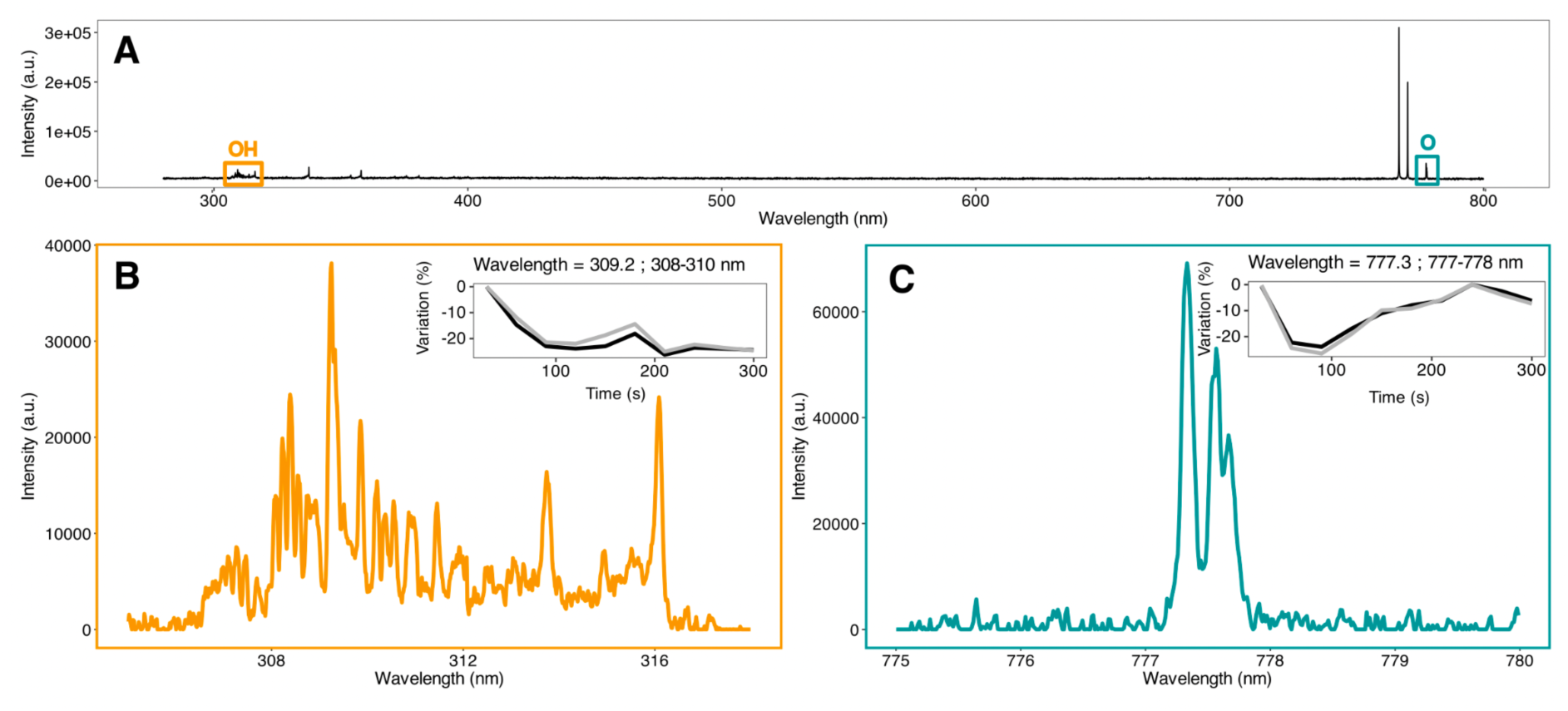
Optical emission spectroscopy. **A.** Optical emission spectrum of the helium-oxygen fed cold atmospheric plasma interacting with the microalgal suspension, acquired between 280 nm and 800 nm; **B.** The highlighted spectral region between 306 nm and 318 nm correspond to the hydroxyl radical •OH(A^2^ Σ^+^→X^2^ π) rovibronic band emissions, but also include the N_2_ second positive system (Table 2); **C.** The highlighted spectral region between 764 nm and 789 nm corresponds to the atomic oxygen O I (3p^5^Pj→3s^5^) emission region.

**Table 2.** Main radicals generated by He/O_2_ cold atmospheric plasma setup. Characterization based on optical emission spectrum (Fig. 2)

| Wavelength (nm) | Compound |
| --- | --- |
| 309.27 | Hydroxyl, $\bullet\text{OH}(A^2\Sigma^+, v' = 0 \rightarrow X^2\Pi, v'' = 0)$ |
| 316.12 | $\text{N}_2$ second positive system, $\text{N}_2(C^3\Pi_u, v' = 1 \rightarrow B^3\Pi_g, v'' = 0)$ |
| 337.35 | $\text{N}_2$ second positive system (0–0) band |
| 357.90 | $\text{N}_2$ second positive system (0–1) band |
| 766.65 and 770.08 | Neutral-potassium K I resonance doublet |
| 777.36; 777.60 and 777.70 | O I triplet ($\text{O I } 3p^5P_J \rightarrow 3s^5S^{\circ}_2$) |

The insets in the upper right corners of panels B and C show variations that represent changes in optical emission intensity and do not constitute direct measurements of species production rates or concentrations. These variations are expressed as % changes relative to the maximum value recorded over the entire experiment. These relatives variations were calculated using two approaches based on: the integrated spectral area over 308–310 nm (*perc_area,* black line), with each value expressed relative to the maximum integrated area recorded throughout the experiment; and the maximum intensity recorded over 308–310 nm (*perc_im,* grey line), with each value expressed relative to the maximum intensity recorded throughout the experiment.

The emission band at 309.27 nm was assigned to the •OH (A^2^Σ^+^, vʹ = 0 → X^2^Π, vʺ = 0) rovibronic system while the one at 316.12 nm was assigned to the N_2_ second positive system, N_2_(C^3^Π_u_, vʹ = 1 → B^3^Πg, vʺ = 0). Additional N_2_ second-positive-system bands were detected at 337.35 and 357.90 nm and assigned to the (0-0) and (0-1) vibrational transitions, respectively. The three closely spaced features near 777 nm (i.e., 777.36; 777.60 and 777.70) were assigned to the O I 3p ^5^P_J → 3s ^5^S°_2_ triplet, whose reference components occur at approximately 777.19, 777.42, and 777.54 nm. In contrast, the features observed at 766.65 and 770.08 nm were assigned to the neutral-potassium K I resonance doublet, rather than to atomic oxygen. During the 300-s observation period, the •OH-associated and O I-associated emission profiles decreased by up to approximately 25.6% and 25.2%, respectively. These changes reflect emission intensity and should not be interpreted directly as changes in radical concentration or production rate.

Infrared thermography revealed a spatially heterogeneous increase in microalgal suspension temperature during cold atmospheric plasma (CAP) exposure. The largest increase occurred near the submerged outlet of the quartz tube, adjacent to the discharge and bubbling region (Supplementary Fig. 1A, B). The mean value constantly increased from approximately 28.5°C at baseline to 32.5°C after 300 s, corresponding to a net increase of approximately 4°C (Supplementary Fig. 1C, D).

In parallel, pH values remained relatively stable during the CAP exposure, from approximately 7.85 at baseline to 7.74 after 5 min. This corresponded to a net change of-0.11 pH unit, or an average change of approximately-0.022 pH unit min^-1^ over the treatment period (Supplementary Fig. 1E).

### Pigment and isoprostanoids profiles of natural microphytobenthic diatom biofilm assemblage dominated by *Pleurosigma strigosum*

The microalgal suspension, derived from the natural microphytobenthic community, reached a final cell concentration of 1.2 × 10^7^ cells per liter. Phototrophic community composition, determined by identifying 1,765 cells, revealed an assemblage of motile benthic diatoms dominated by *Pleurosigma strigosum* accounting for 51.3%. The remaining composition was shared among other diatom taxa, including *Navicula sp*., *Gyrosigma sp*., *Entomoneis sp*., and other *Pleurosigma sp*. (Supplementary Fig. 2). Occasional meiofaunal individuals, including *Nematoda* and *Arthropoda* (e.g., copepods and nauplius larvae), were also observed in the suspension.

A total of 65 metabolites were detected and identified, 53 of which were lipophilic pigments. The lipophilic pigment profile of the diatom assemblage was largely dominated by chlorophyll *a* and its derivatives, fucoxanthin, and chlorophyll *c_2_*, accounting for 58.3%, 25.1%, and 6.2% of the average relative abundance, respectively. The remaining composition included neoxanthin, unidentified carotenoids, pheopigments, the diadinoxanthin + diatoxanthin pool, β-carotenes, and lutein (2.5%, 2.4%, 2.1%, 2%, 1.5%, and 0.1%, respectively; Fig. 3).

**Figure 3.**
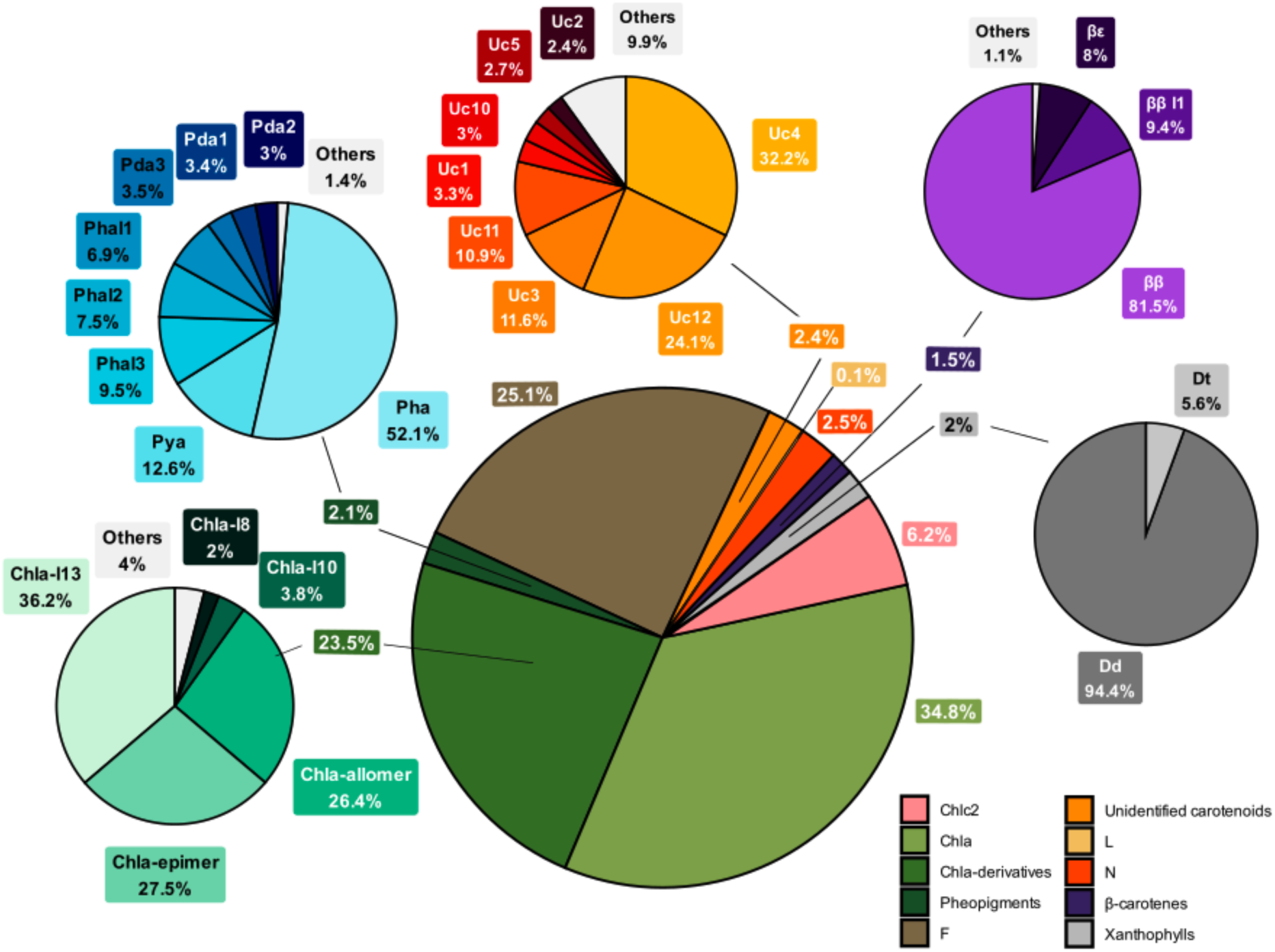
Average composition of the lipophilic pigment profile. Average lipophilic pigment composition (%, calculated on pigment concentration) across samples (n = 56). The “Others” category includes lipophilic pigments with < 2% relative abundances of the group’s pool. The abbreviations for pigment names are defined in Supplementary Table 2.

Among the 54 available oxylipin standards derived from ω3 and ω6 polyunsaturated fatty acid, profiling identified 12 individual compounds (8 excluding stereoisomeric duplicates) belonging to three major classes: isoprostanes, neuroprostanes, and phytoprostanes. These isoprostanoids were derived from four precursor polyunsaturated fatty acids: eicosapentaenoic acid (C20:5 ω3), docosahexaenoic acid (C22:6 ω3), α-linolenic acid (C18:3 ω3), and arachidonic acid (C20:4 ω6). The isoprostanoid profile of the diatom assemblage was largely dominated by eicosapentaenoic acid-derived isoprostanes, which accounted for 68.8% of the total relative isoprostanoid pool. The remaining composition consisted of docosahexaenoic acid-derived neuroprostanes (15.6%), α-linolenic acid-derived phytoprostanes (11.4%), and arachidonic acid-derived isoprostanes (4.2%) (Fig. 4). At the individual metabolite level, the eicosapentaenoic acid-derived stereoisomers 5-*epi*-5-F_3t_-IsoP and 5-F_3t_-IsoP were the most abundant isoprostanoids detected. Notably, the stereoisomer 5-*epi*-5-F_3t_-IsoP represented more than 70% of the total 5-F_3t_-IsoP stereoisomer pool.

**Figure 4.**
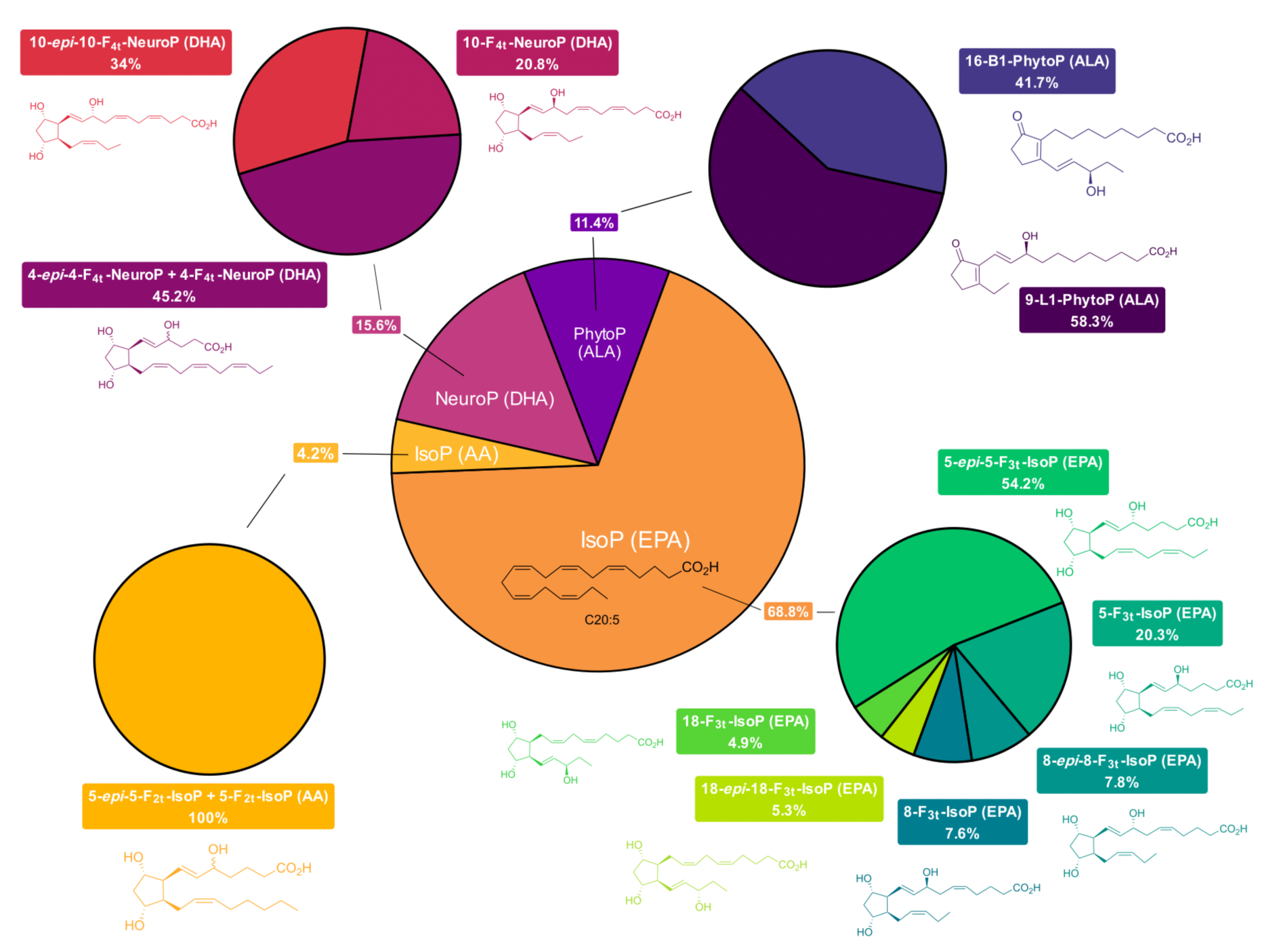
Average composition of the isoprostanoid profile. Average isoprostanoids composition (%, calculated on raw chromatogram area) across samples (n = 56). Eicosapentaenoic acid, EPA; Docosahexaenoic acid, DHA; ɑ-linolenic acid, ALA; Arachidonic acid, AA; Isoprostanes, IsoP; Neuroprostanes, NeuroP; Phytoprostanes, PhytoP.

### Differential cold atmospheric plasma effects on diatom pigment and isoprostanoid compositions over time

The impact of oxidative conditions induced by CAP treatments on the diatom assemblage was investigated through pigment and isoprostanoid compositional responses. Based on these compositions, samples showed no clear clustering within the MFA space under the 2-min CAP treatment (Supplementary Fig. 3A, B). Among the samples treated with 2-min CAP, only those collected immediately after CAP exposure formed a distinct cluster, which was apparent only in the BC-MFA representation, whereas samples collected after 30 and 180 min of recovery grouped with the controls (Supplementary Fig. 3C, D). However, this treatment-related structuring was not statistically significant (Monte-Carlo test; *p* > 0.05).

In contrast, the 5-min CAP treatment produced clear clustering within the MFA space (Fig. 5A, B), with the immediate post-exposure and 30-min recovery samples forming distinct clusters, whereas the 180-min recovery samples grouped with the controls. This treatment-related structuring accounted for 28.7% of the total explained inertia (first two BC-MFA dimensions Monte Carlo test; *p* < 0.05; Fig. 5C, D).

**Figure 5.**
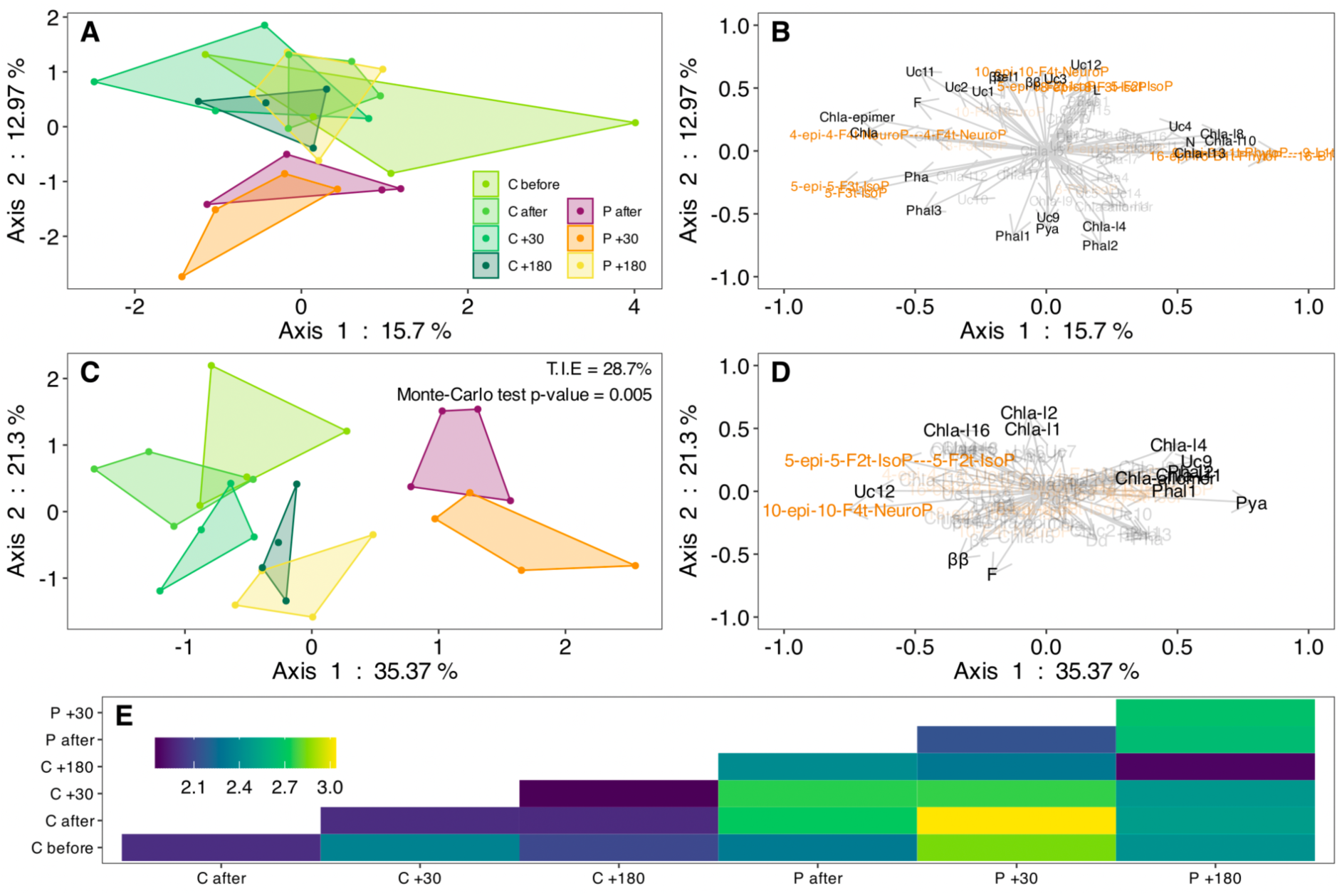
Lipophilic pigment and isoprostanoid responses to 5-min cold atmospheric plasma. **A.** Projection of samples in the factorial space derived from a Multiple Factor Analysis (MFA) based on their lipophilic pigment and isosprostanoid composition (%) across treatments; **B.** Projection of variable vectors in the MFA factorial space, showing only variables with cos^2^ > 0.2 for improved readability. Vectors directions and lengths indicate their correlations and relative contributions to the structuring of the factorial space; **C.** Between-class analysis performed on the MFA object (BC-MFA), constrained by the explanatory treatment variable. The total inertia explained (T.I.E.) reflects the proportion of variance associated with the partition imposed by the explanatory variable; **D.** Projection of variable vectors in the BC-MFA factorial space, showing only variables with cos^2^ > 0.2 for improved readability. Vectors directions and lengths indicate their correlations and relative contributions to the structuring of the factorial space; **E.** Heatmap of Euclidean distances between group centroids performed across all MFA dimensions. 5-min cold atmospheric plasma treatments (n = 4): Control before, C before; Control after, C after; Control +30 min after treatment cessation, C +30; Control +180 min after treatment cessation, C +180; Cold atmospheric plasma treated-samples after, P after; Cold atmospheric plasma treated-samples +30 min after treatment cessation, P +30; Cold atmospheric plasma treated-samples +180 min after treatment cessation, P +180.

The first BC-MFA axis primarily reflected the impact of the treatments, whereas the second axis captured the temporal trajectory of the samples (Fig. 5C).

The representation of variable groups further indicated that, for both CAP treatment durations, isoprostanoids and pigments contributed > 50 % to the structuring of axis 1 (temporal trajectory) of the multivariate MFA spaces (Fig. 5A), with isoprostanoids consistently showing a greater contribution than pigments (Supplementary Fig. 4). In contrast, axis 2 of the MFA (treatments) was almost exclusively structured by pigments under both CAP treatment durations, with their contribution reaching up to 80%.

The two-way PERMANOVA performed on the global pigment dataset confirms the structuring of pigment compositions observed in the multivariate space and reveals significant effects of time and CAP treatments on pigment composition (Two-way PERMANOVA; *p* < 0.05 for both factors; *R^2^* _temps_ = 0.19; *R^2^* _treatment_ = 0.43), whereas their interaction was not significant (Two-way PERMANOVA; *p* > 0.05). As multivariate dispersions were homogeneous among groups (multivariate dispersion test; *p* > 0.05), the observed differences reflect compositional changes rather than differences in within-group dispersion.

In contrast, the two-way PERMANOVA performed on the entire isoprostanoid dataset revealed a significant effect of CAP treatments on isoprostanoid composition (Two-way PERMANOVA; *p* = 0.002; *R^2^* _treatment_ = 0.18), but no significant effect of time or of the interaction between the two factors (Two-way PERMANOVA; *p* > 0.05). This result should nevertheless be interpreted with caution, as multivariate dispersions differed significantly among groups (multivariate dispersion test; *p* = 0.007), indicating that the effect associated with CAP treatments may partly reflect differences in within-group dispersion.

Euclidean distances between the centroids of the 5-min CAP treatment groups, across all MFA dimensions, revealed that sample collected immediately after and +30 min after cessation of CAP treatment were the two groups showing the greatest dissimilarity from the controls, with the +30 min samples being the most dissimilar (Fig. 5E). Hierarchical clustering of these centroid-based distances further highlighted both treatment-related differences and the temporal relationships between sample groups (Supplementary Fig. 5). A first clustercomprised the samples collected immediately after and +30 min after cessation of the 5-min CAP treatment, consistent with persistent CAP-induced alterations in their integrated pigment and isoprostanoid profiles. In contrast, the remaining treatment groups formed a second cluster that primarily reflected the natural temporal remodelling in metabolic profiles throughout the experiment. Within this cluster, the control treatments formed a distinct subcluster, while samples collected +180 min after CAP treatment cessation remained relatively close to the controls but distinct from them.

At the individual metabolite level, some lipophilic pigments and isoprostanoids showed response dynamics that may help explain the significant clustering observed according to time and under the 5-min CAP treatment, with distinct response dynamics across compounds (Fig. 6).

**Figure 6.**
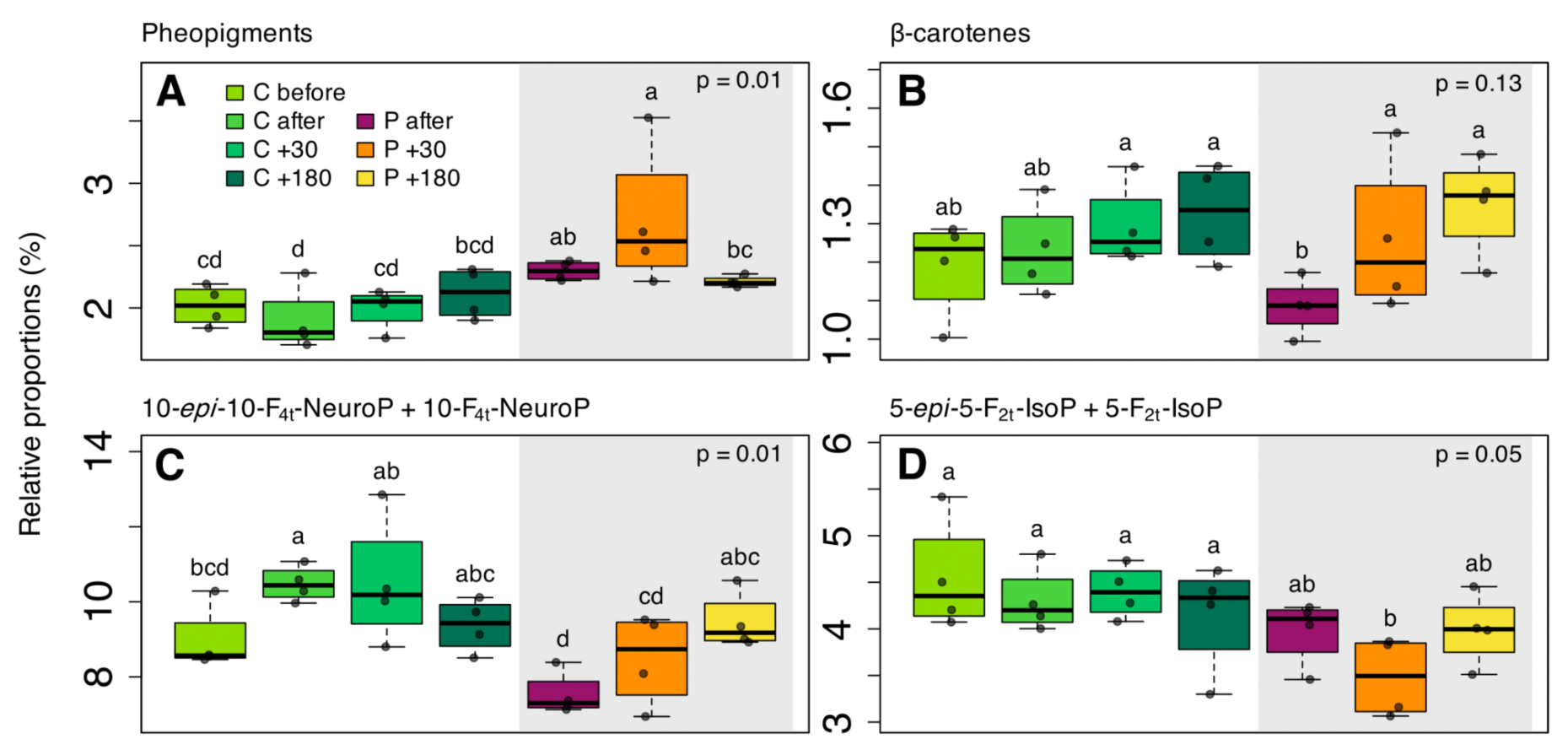
Pigment group and individual isoprostanoid relative proportion variations under 5-min cold atmospheric plasma. Boxplots showing the relative proportions (%) of the most relevant compounds driving diatom metabolic responses under 5-min cold atmospheric plasma treatments; **A.** pheopigments group, including pheophytin *a*, pheophorbide *a* and pyropheophytin *a* forms; **B.** β-carotenes group, including β-β-carotene and β-દ-carotene forms; **C.** 10-F_4t_-neuroP group, including 10-*epi*-10-F_4t_-neuroP and 10-F_4t_-neuroP stereoisomers; **D.** 5-F_2t_-isoP group, including 5-*epi*-5-F_2t_-isoP and 15-F_2t_-isoP stereoisomers. Relative proportions of lipophilic pigments were calculated on pigment concentrations. Relative proportions of isoprostanoids were calculated on raw chromatogram areas. 5-min cold atmospheric plasma treatments: Control before, C before; Control after, C after; Control +30 min after treatment cessation, C +30; Control +180 min after treatment cessation, C +180; Cold atmospheric plasma treated-samples after, P after; Cold atmospheric plasma treated-samples +30 min after treatment cessation, P +30; Cold atmospheric plasma treated-samples +180 min after treatment cessation, P +180. Cold atmospheric plasma treatment groups were statistically tested using the Van der Waerden test (n = 4).

The pheopigment group was among the most responsive metabolite groups, showing a post-treatment increase in their relative proportions that persisted during the post-treatment period and reached a maximum at +30 min after cessation of the 5-min CAP treatment compared with controls, before returning to comparable control levels +180 min after treatment (Van der Waerden test; *p* < 0.05; Fig. 6A). This overall group-level response, mainly driven by pheophytin *a*, pheophytin *a*-like 1 and 2, as well as pyropheophytin *a* which showed the clearest response, masked the dynamics of pheophorbides *a* 1 and 2, which exhibited an opposite trend under CAP treatment (Supplementary Fig. 6). In contrast to the treatment effect, pheopigments did not appear to be substantially affected by time, based on the temporal evolution observed in the controls. The β-carotenes also showed a tendency towards a post-treatment response, predominantly driven by β-β-carotene form, but with a distinct response dynamic, as their relative proportions initially decreased and then progressively returned to comparable control levels after 180 min (Van der Waerden test; *p* = 0.13; Fig. 6B and Supplementary Fig. 6). This treatment-related response also appeared to be accompanied by a temporal component, based on the evolution observed in the control samples. Other pigments, such as fucoxanthin, also showed changes in their relative proportions over time, but did not appear to be substantially affected by the 5-min CAP treatment (Supplementary Fig. 8). Finally, the diadinoxanthin-diatoxanthin antioxidant xanthophyll cycle showed no significant diadinoxanthin de-epoxidation state variation under any CAP treatment compared with control samples (Supplementary Fig. 6).

Regarding isoprostanoids, only two compounds showed significant changes in their relative proportions following the 5-min CAP treatment: 10-F_4t_-neuroP, derived from docosahexaenoic acid, and 5-F_2t_-IsoP, derived from arachidonic acid (Fig. 6C, D). The neuroprostane 10-F_4t_-neuroP showed a post-treatment decrease in relative proportions, followed by a progressive return to comparable control levels +180 min after cessation of CAP treatment, a dynamic mainly driven by the 10-*epi*-10-F_4t_-neuroP form (Supplementary Fig. 7). Similarly, 5-F_2t_-IsoP showed a post-treatment decrease in relative proportions that persisted throughout the post-treatment period and reached a minimum +30 min after cessation of CAP treatment, before returning to comparable control levels +180 min after cessation. No isoprostanoid derived from the dominant eicosapentaenoic acid precursor showed a detectable response to the 5-min CAP treatment.

### Cold atmospheric plasma causes only minor and transient changes in diatom photosynthetic performance

Despite the significant impact of the 5-min CAP treatment on lipophilic pigments and isoprostanoid profiles, the diatom assemblage maintained its photosynthetic performances, with most photophysiological parameters remaining unaffected, although a small but statistically significant transient reduction in Fv/Fm occurred (Van der Waerden test; *p* > 0.05; Supplementary Table 3 and Supplementary Fig. 8). The diatoms exhibited very high maximum quantum efficiencies of PSII Fv/Fm (> 0.60), which nevertheless showed a slight decrease following treatment cessation under both the 2-min and 5-min CAP treatments, by an average of-0.02 and-0.03 units, respectively (Van der Waerden test; *p* < 0.05; Fig. 7). In both cases, this decrease was followed by a progressive post-treatment recovery towards control values, although values remained slightly different between controls and samples collected +180 min after cessation of CAP treatment. This dynamic was nevertheless consistent with the progressive convergence observed in pigment and isoprostanoid profiles over time following treatment cessation.

**Figure 7.**
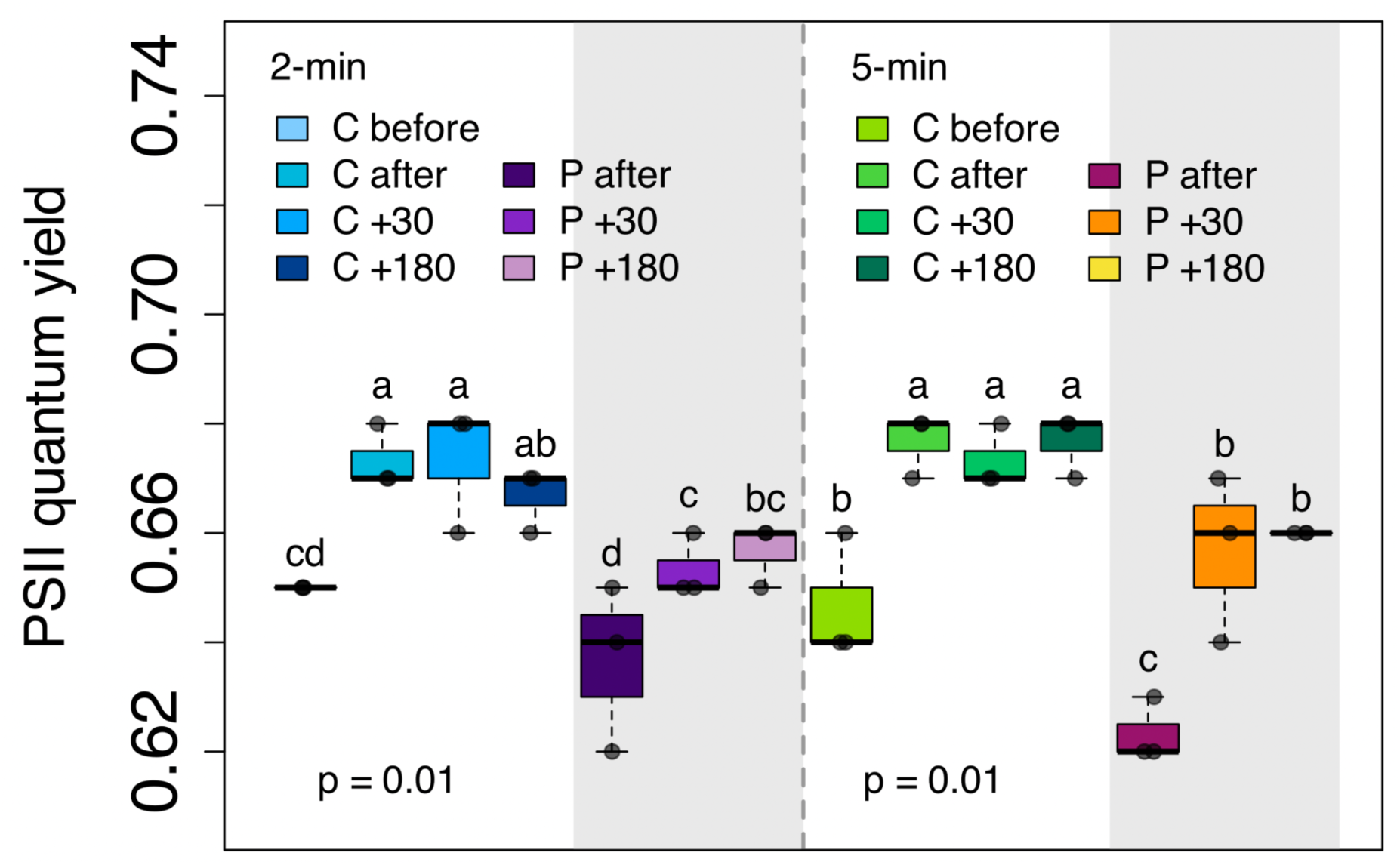
Maximum quantum efficiencies of PSII (Fv/Fm) photosynthetic parameter. Boxplots showing the changes in the maximum quantum efficiencies of PSII (Fv/Fm) photosynthetic parameter under 2-min and 5-min cold atmospheric plasma treatments: Control before, C before; Control after, C after; Control +30 min after treatment cessation, C +30; Control +180 min after treatment cessation, C +180; Cold atmospheric plasma treated-samples after, P after; Cold atmospheric plasma treated-samples +30 min after treatment cessation, P +30; Cold atmospheric plasma treated-samples +180 min after treatment cessation, P +180. Cold atmospheric plasma treatment groups were statistically tested using the Van der Waerden test (n = 4).

### No detectable loss of diatom membrane integrity following 5-min cold atmospheric plasma treatment

SYTOX Green fluorescence was used to assess cell membrane integrity, as the dye penetrates cells with compromised plasma membranes and binds to nucleic acids, resulting in green nuclear fluorescence.

Positive controls showed strong nuclear fluorescence, confirming the ability of the dye to enter membrane-compromised cells, whereas negative controls showed little to no nuclear fluorescence, consistent with the exclusion of the dye by intact cell membranes (Fig. 8A, B). At +180 min after cessation of the 5-min CAP treatment, the diatom assemblage showed no marked nuclear fluorescence compared with the negative control and differed substantially from the strong nuclear fluorescence observed in the positive control (Fig. 8C). Residual autofluorescence was also observed in extracellular polymeric substances aggregates and the sediment substrata, potentially reflecting fluorescence associated with extracellular material, including extracellular DNA.

**Figure 8.**
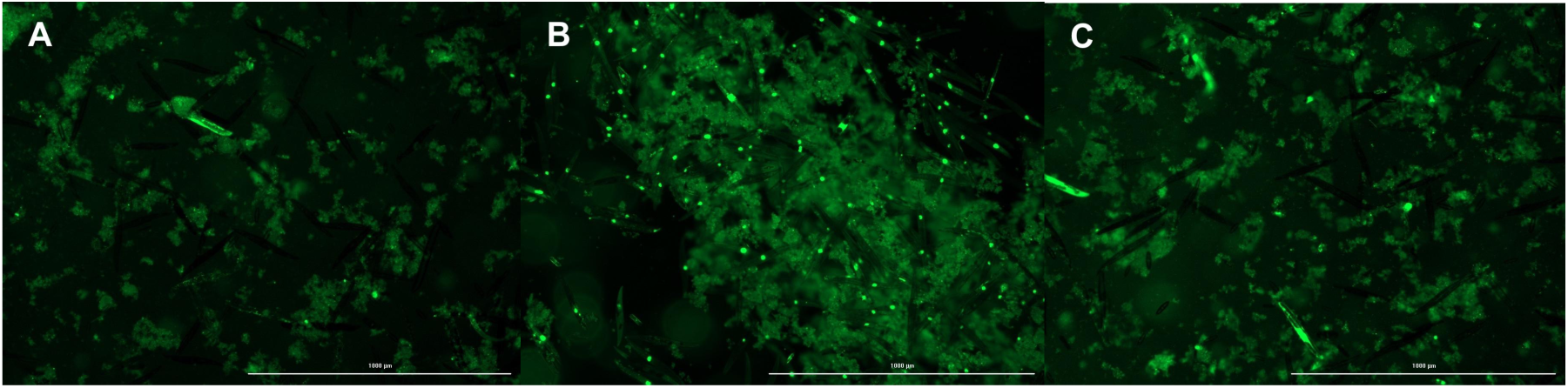
Diatom viability following 5-min cold atmospheric plasma treatment. Representative fluorescence microscopy images of SYTOX Green fluorescence acquired in GFP mode (λ _LED excitation_ = 469 nm and λ _LED emission_ = 525 nm), +180 min after cold atmospheric plasma treatment cessation, showing: **A.** the negative control; **B.** the positive control; **C.** the 5-min cold atmospheric plasma treated sample. No green fluorescence was detected in the filtered seawater blank.

## Discussion

### Impact of cold atmospheric plasma on a natural diatom assemblage dominated by *Pleurosigma strigosum*

#### Characterization of the natural diatom assemblage and cold atmospheric plasma-induced oxidative conditions

The natural microphytobenthic assemblage investigated here was composed predominantly of motile epipelic benthic diatoms, with *Pleurosigma strigosum* as the dominant taxon (Supplementary Fig. 2), a characteristic structure of diatom-dominated communities inhabiting comparable muddy sediment substrata (Underwood & Barnett, 2006; Méléder et al., 2007), and consistent with previous metabarcoding-and microscopy-based characterizations of motile communities from the same study site (Gaubert-Boussarie et al., 2020; Doose & Hubas, 2024; Desparmet et al., 2026a). The dominance of the large *P. strigosum* cells likely resulted in a disproportionately high contribution to the phototrophic biovolume of the assemblage, such that the molecular and physiological responses measured here are expected to be largely driven by this taxon. This interpretation is further supported by the pigment profile, which was characterized by high relative abundances of chlorophyll *a*, fucoxanthin, and chlorophyll *c2*, together with xanthophylls such as diadinoxanthin and diatoxanthin (Fig. 3), a chemotaxonomic signature consistent with diatoms and closely matching profiles previously reported for similar microphytobenthic assemblages (Roy et al., 2011; Desparmet et al., 2026a; Desparmet et al., 2026b). Likewise, the isoprostanoid profile was dominated by polyunsaturated fatty acid-derived compounds originating from ω3 fatty acids, with eicosapentaenoic acid-derived isoprostanes accounting for the largest fraction (Fig. 4), as expected from the predominance of eicosapentaenoic acid in the polyunsaturated fatty acid pool typically reported in diatoms (Dunstan et al., 1993; Dijkman et al., 2010), and consistent with previous characterization of this *P. strigosum*-dominated microphytobenthic assemblage (Desparmet et al., 2026b).

Considering the high Fv/Fm values (Fig. 7; Prins et al., 2020; Morelle et al., 2025a; Morelle et al., 2026), together with microscopic identifications, pigmentary compositions, and isoprostanoid profiles, the results confirm that the experimental model was in good physiological condition prior to treatment and representative of a natural epipelic diatom assemblage dominated in biovolume by *P. strigosum*.

Cold atmospheric plasma (CAP) treatment induces environmental changes primarily characterized by the generation of a substantial and complex mixture of reactive species (Fig. 2 and Table 2; Perinban et al., 2019; Mai-Prochnow et al., 2021; Thompson et al., 2025). Since the concomitant temperature and pH changes were moderate compared with environmental fluctuations that benthic diatoms may encounter in intertidal environments (Supplementary Fig. 3; Serôdio, 2021), they are unlikely to have been the primary factors of the observed biological responses. Here, the bubbling experimental setup resulted in exposure to an exogenous oxidative environment induced by the CAP, maximising continuous contact between plasma-generated reactive species and the natural diatom assemblage (Fig. 1A). OES demonstrated OH-associated and atomic-oxygen emission from the plasma discharge (Fig. 2). Additional reactive species, including ^1^O_2_, O_2_^•-^ and longer-lived species such as H_2_O_2_, may be generated through CAP–liquid chemistry, as reported for comparable systems (Perinban et al., 2019; Mai-Prochnow et al., 2021; Thompson et al., 2025); however, these species were not individually quantified here. ROS production and accumulation in plasma-activated water strongly depend on several parameters, including power supply, voltage, frequency, gas mixture, and treatment duration (Mai-Prochnow et al., 2021). The diffusion of reactive species from the CAP into the aqueous phase involves numerous simultaneous physical and chemical processes rather than a simple transfer from one compartment to another (Perinban et al., 2019; Mai-Prochnow et al., 2021). Depending on the CAP system, a fraction of the short-lived species (ns to s) generated will act as transient reactive intermediates that are rapidly consumed or converted into more stable species, such as the long-lived H_2_O_2_, which can accumulate in plasma-activated water, contributing to its sustained oxidative activity and allowing reactive chemistry to diffuse into the aqueous phase. However, when short-lived species are generated sufficiently close to biological targets, as may occur in bubbling CAP setup, they may also react directly with nearby biological substrates before being transformed, thereby contributing to local oxidative stress without necessarily accumulating in the aqueous microalgal suspension. In addition, exogenous long-lived species induced by CAP can react with extracellular polymeric substances and other biological components, leading to the local formation of short-lived reactive species (Perinban et al., 2019; Mai-Prochnow et al., 2021; Thompson et al., 2025).

#### Cold atmospheric plasma-induced oxidative condition transiently alters pigment profiles without compromising diatom photophysiology performances

Despite an oxidative exposure associated with physicochemical changes, MFA and BC-MFA analyses highlighted limited changes in pigment and isoprostanoid profiles, which were dependent on CAP treatment duration, with a significant overall effect identified only for the 5-min CAP treatment (Figs. 5 and 6). Together with the overall stability of photophysiological parameters, the high post-treatment Fv/Fm values (Fig. 7) and the maintenance of membrane integrity further indicate that the 5-min CAP treatment did not induce detectable excess cell mortality (Fig. 8), these results suggest that the treatments remained within the physiological tolerance range of the assemblage, with no major detectable loss of membrane integrity or photosynthetic function.

Multivariate space structuring, together with PERMANOVA results (Figs. 5, supplementary s3–5), highlighted significant and additive effects of treatment and temporal components on pigment profiles, with no significant interaction. In contrast, isoprostanoid profiles appeared less responsive, showing only a weaker but significant effect of CAP treatment, although this result should be interpreted with caution due to heterogeneous multivariate dispersions among groups. Overall, these results indicate that the response to CAP-induced oxidative conditions was predominantly structured by pigments rather than lipid-related metabolites, consistent with their similarly predominant contribution to metabolic remodelling under different photooxidative treatments (Desparmet et al., 2026b).

The metabolic changes observed over time in control samples likely reflect endogenous cellular regulation, as epipelic diatoms are known to tightly regulate multiple biological processes in anticipation of predictable environmental variations, such as diel and tidal cycles, thereby conferring a survival advantage through anticipatory adjustment (Consalvey et al., 2004; Coelho et al., 2011; Annunziata et al., 2019; Manzotti et al., 2025). Finally, the progressive convergence of Fv/Fm, as well as pigment and isoprostanoid profiles in treated samples, toward those of controls over time following cessation of CAP treatments, suggests that the plasma-induced metabolic changes are partially reversible, potentially reflecting a degree of metabolic resilience to transient oxidative perturbation.

The deleterious impact of plasma-induced oxidative conditions on diatoms is highlighted by the increase in pheopigments following CAP treatment, with this effect persisting for at least +30 min after treatment cessation. The formation of these compounds occurs through the catabolic degradation of chlorophyll *a*, whereby Mg^2+^ removal first leads to the formation of pheophytin *a,* followed by the loss of the phytol chain to form pheophorbide *a* (Queiroz Zepka et al., 2019). In contrast, pyropheophytin *a* results from the loss of the methoxycarbonyl group from pheophytin *a* under specific conditions, particularly thermal conditions (Schwartz & Von Elbe, 1983). In parallel, the decrease in β-carotenes levels may suggest its mobilization in diatoms in response to oxidative conditions, given its antioxidant properties through direct radical scavenging, in addition to its role as a precursor in the biosynthesis of other pigments, including xanthophylls and fucoxanthin (Rijstenbil, 2003; Kuczynska et al., 2015). In contrast to photooxidative stress, these plasma-induced oxidative conditions altered pigment profiles without inducing detectable changes in the diadinoxanthin-diatoxanthin cycle or a decrease in photosynthetic performance, a response profile previously observed by Desparmet et al. (2026a). The progressive return of the plasma-modified pigment profiles toward control profiles suggests that diatoms may actively manage the chlorophyll *a* degradation products generated, while progressively restoring their relative β-carotenes abundance following cessation of CAP exposure.

#### Isoprostanoid of microphytobenthic diatom assemblage under cold atmospheric plasma treatment vs photooxidative stress

Due to the targeted subsampling of the community surface layer, fewer isoprostanoids were identified here than reported in microphytobenthic sediment samples by Doose et al. (2024), who detected 26 isoprostanoids (17 when excluding stereoisomeric duplicates). Nevertheless, the more restricted community analyzed here may provide a more specific representation of diatom-associated isoprostanoids signatures, whereas the broader sediment samples may include isoprostanoids originating from abundant meiofauna, including nematodes. Despite some differences, the overall isoprostanoid profiles characterized in Doose et al. (2024) were remarkably similar, with a strong predominance of eicosapentaenoic acid-derived 5-F_3t_-isoP, particularly the stereoisomer 5-*epi*-5-F_3t_-isoP, which appears to be preferentially formed in diatoms, possibly due to a specific radical-mediated reaction favored by the conformational accessibility of the precursor, together with other factors operating *in vivo*.

The absence of a response in the major eicosapentaenoic acid-derived isoprostanoids, despite concomitant chlorophyll *a* degradation, provides further insight into the exogenous, untargeted, and relatively mild nature of the oxidative conditions induced here by CAP treatments, in contrast to the photooxidative stress applied by Doose et al. (2024), which is known to generate endogenous chloroplastic ROS (Mizrachi et al., 2019; Foyer & Hanke, 2022), thereby promoting the formation of eicosapentaenoic acid-derived isoprostanoids. Moreover, isoprostanoid profiles differed depending on the intensity of photooxidative stress, suggesting that the isoprostanoid response of diatoms may not be purely stochastic under specific conditions (Doose et al., 2024).

All these results raise the following spatial and mechanistic question: assuming that exogenously generated CAP ROS induce oxidative damage primarily through the direct and non-specific oxidation of the nearest cellular targets, how can they induce changes in pigment ratios (Fig. 6A, B), particularly an increase in chlorophyll *a* degradation products, without causing a concomitant increase in eicosapentaenoic acid-derived isoprostanoids? This is particularly intriguing given that most chlorophyll *a* is localized within the thylakoid membranes, where it is associated with photosynthetic complexes, whereas EPA is also highly abundant in thylakoid membrane glycolipids, particularly MGDG and DGDG, while being more broadly distributed across cellular compartments (Büchel et al., 2022).

This suggests that the effects induced by plasma-generated ROS may not simply result from the direct, non-specific oxidation of the most abundant polyunsaturated fatty acid substrates. Two non-exclusive mechanisms could account for this apparent discrepancy. First, oxidation may be strongly constrained by cellular architecture, with precursor physicochemical properties, incorporation into specific lipid classes, subcellular localization relative to endogenous or exogenous ROS sources, and local ROS concentrations collectively determining their susceptibility to oxidation (Jahn et al., 2008; Galano et al., 2017; Ahmed et al., 2020). Second, and more speculatively, diatoms may preferentially preserve EPA-rich structural lipids to maintain membrane integrity, while oxidative pressure is borne by other polyunsaturated fatty acid pools or modulated through regulated lipid remodelling, potentially contributing to the formation of isoprostanoids derived from less abundant precursors and to non-random oxidation signatures. Such a mechanism could potentially contribute to explaining why the dynamics of isoprostanoids appear to be non-stochastic across different photooxidative intensities as observed by Doose et al. (2024), and may therefore represent an interesting avenue for further investigation.

Although the biological roles of isoprostanoids remain largely unexplored, if isoprostanoid signatures are modulated according to the intensity of a given stress, they could serve as biomarkers of oxidative stress, providing cells with valuable information about their environmental conditions, the physiological state of neighboring cells, and their own physiological status (Bultel-Poncé et al., 2016; Galano et al., 2017; Doose et al., 2024). This possibility is particularly relevant given that diatoms are known to perceive and integrate oxidative signals to trigger metabolic, transcriptional, and behavioural functional responses (Valle et al., 2014; Carrara et al., 2021; Pivato et al., 2023; Flori et al., 2025; Desparmet et al., 2026a).

#### Isoprostanoid and enzymatic oxylipin roles across living systems

In this experimental setup, only the minor isoprostanoids 10-F_4t_-neuroP and 5-F_2t_-IsoP showed significant changes. In addition to their role as biomarkers under various oxidative conditions, including H_2_O_2_ exposure, copper-induced stress, and increased irradiance (Lupette et al., 2018; Vigor et al., 2020; Doose et al., 2024), some isoprostanes have been reported to exert anti-inflammatory effects, induce vasoconstriction and apoptosis, and modulate neurotransmitter release, whereas, some neuroprostanes have been associated with anti-inflammatory, anti-arrhythmic, cardioprotective, and antiproliferative activities. Moreover, phytoprostanes have been linked to neuroprotective effects, the promotion of cellular differentiation, the stimulation of antimicrobial secondary metabolite production which may enhance tolerance to biotic stress, and have also been shown to activate plant defence and detoxification responses by inducing the expression of stress defence-related genes (Bultel-Poncé et al., 2016; Galano et al., 2017; Vigor et al., 2018; Knieper et al., 2023). Interestingly, some isoprostanoids may also act as signaling molecules even in the absence of the enzymes required for the biosynthesis of related enzymatic oxylipins. This is notably the case in *P. tricornutum*, in which cyclopentane isoprostanoids have been shown to induce cellular responses despite the absence of any identified canonical enzymatic machinery for cyclopentane oxylipin biosynthesis (Lupette et al., 2018). In addition, some isoprostanoids and enzymatically derived oxylipins may converge on common cellular responses, exhibiting functional overlap and engaging shared signaling pathways, while not being strictly equivalent (Knieper et al., 2023). This phenomenon is notably illustrated by 15-F_2t_-IsoP, an isoprostanoid primarily generated through non-enzymatic free-radical-mediated peroxidation, which can elicit functional responses through activation of the thromboxane TP receptor (TPR), for which it acts as a partial agonist (Bauer et al., 2014). Taken together, this growing body of evidence further supports the importance of isoprostanoids as biologically relevant signals that can be interpreted by cells to modulate meaningful biological responses, similarly to enzymatically derived oxylipins.

Here, the significant changes in isoprostanoids reflect modifications in exogen conditions induced by CAP treatments. Unfortunately, the biological significance of the observed changes in 10-F_4t_-neuroP and 5-F_2t_-IsoP remains unclear, as no specific direct biological roles or associated cellular responses beyond their use as biomarkers of oxidative damage have yet been established (Galano et al., 2017; Ahmed et al., 2020; Linares-Maurizi et al., 2023). However, the decrease in their relative proportions with increasing oxidative pressure may appear counterintuitive. Vigor et al. (2020) previously reported decreases in isoprostanoids following H_2_O_2_-induced oxidative stress in *P. tricornutum*. Under oxidative conditions, such decreases could potentially be explained, at least in part, by further β-oxidation and reduction processes, whereby 10-F_4t_-neuroP may undergo additional metabolic transformations leading to the formation of more stable end-products, as hypothesized by Ahmed et al. (2020) and demonstrated for 7-F_4t_-neuroP (Lawson et al., 2006).

## Conclusion

The present work investigated changes in pigment and isoprostanoid profiles, as well as photosynthetic performance, in response to bubbling cold atmospheric plasma treatments, mimicking photooxidative stress without light exposure, in a motile diatom assemblage dominated by *Pleurosigma strigosum* and originating from a natural microphytobenthic community. Our results indicate that pigment and isoprostanoid profiles can reveal changes in environmental oxidative conditions before detectable alterations in photosynthetic performance occur, and may therefore provide complementary signatures of oxidative environmental perturbations. In fact, cold atmospheric plasma induced physicochemical changes characterized by an oxidative challenge, leading to selective and transient metabolic perturbations, primarily reflected in lipophilic pigment profiles and, to a lesser extent, in isoprostanoid profiles, while largely preserving photosynthetic performance. The extent of these perturbations depended on treatment duration, with weaker effects after 2 min than 5 min. The greater divergence from control profiles peaked at +30 min after the 5-min treatment cessation and progressively converged toward control levels by +180 min after treatment cessation. These largely reversible changes were particularly characterized by a transient increase in the relative proportions of pheopigments, accompanied by transient decreases in β-carotenes, 10-F_4t_-neuroP and 5-F_2t_-IsoP relative proportions.

This study provides new insights into how oxidative environmental perturbations affect marine phototrophic microbiome, enhancing our understanding of the ecological success of diatoms in highly variable environments, including intertidal mudflats, where oxidative conditions can fluctuate rapidly. Further in-depth investigation into the *in vivo* formation of isoprostanoids, considering the abundance and intrinsic physicochemical properties of their precursors, as well as their cellular localization and proximity to ROS sources, should help clarify the extent to which isoprostanoid production is driven by stochastic, spatially constrained oxidative processes and whether specific underlying cellular mechanisms may also contribute to shaping their formation.

## Data availability

All scripts and data presented in this article and used to generate the figures and perform the analyses are available on GitHub (https://github.com/adesparmet/Plasma-Isoprostanoid-Pigment-Diatoms).

## Author contributions

**AD** (Conceptualization, Methodology, Investigation, Data curation, Formal analysis, Visualization, Writing – original draft); **CO** (Data curation, Validation, Resources, Supervision, Writing – review & editing); **CV** (Validation, Resources, Writing – review & editing); **TD** (Validation, Resources); **GR** (Data curation, Resources);

**VG** (Data curation, Resources); **TD** (Conceptualization, Methodology, Data curation, Validation, Resources, Supervision, Writing – review & editing); and **CH** (Conceptualization, Formal analysis, Validation, Supervision, Writing – review & editing).

## Supplementary figures and tables

**Figure s1.**
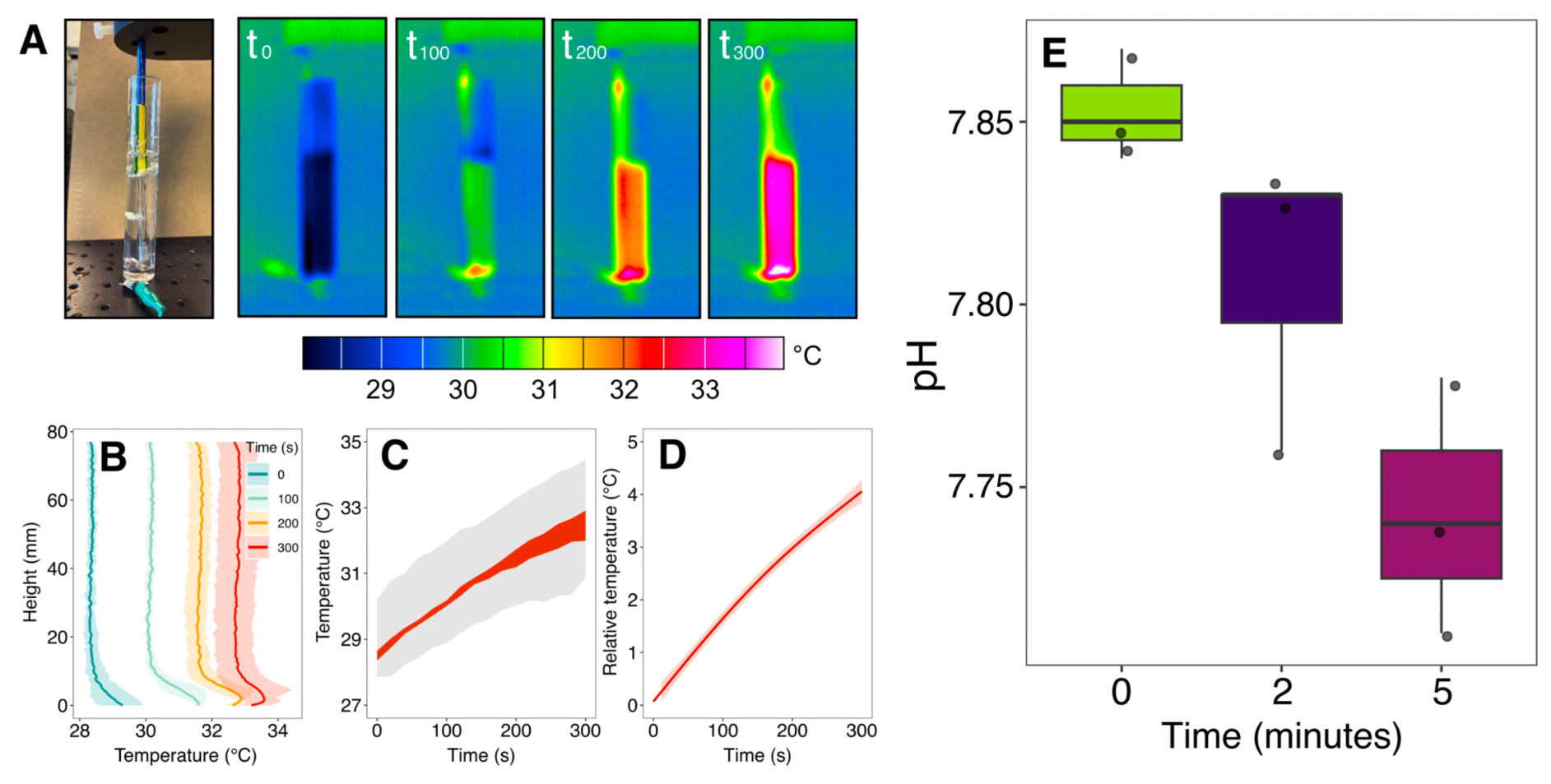
Microalgal suspension physicochemical changes under cold atmospheric plasma. **A.** Infrared thermographic photograph of the quartz tube immersed into the microalgal suspension, followed by four representative IR images presented chronologically from left to right and acquired at successive stages of cold atmospheric plasma treatment, to show the evolution of the microalgal suspension temperature (°C); **B.** Axial profiles of the microalgal suspension temperature at 0, 100, 200, and 300 of cold atmospheric plasma exposure. The shaded regions indicate the variability associated with each profile; **C and D.** Temporal evolution of absolute and relative mean temperature increases of the microalgal suspension; **E.** Boxplots showing pH (unitless) evolution in the microalgal suspension during cold atmospheric plasma treatment.

**Figure s2.**
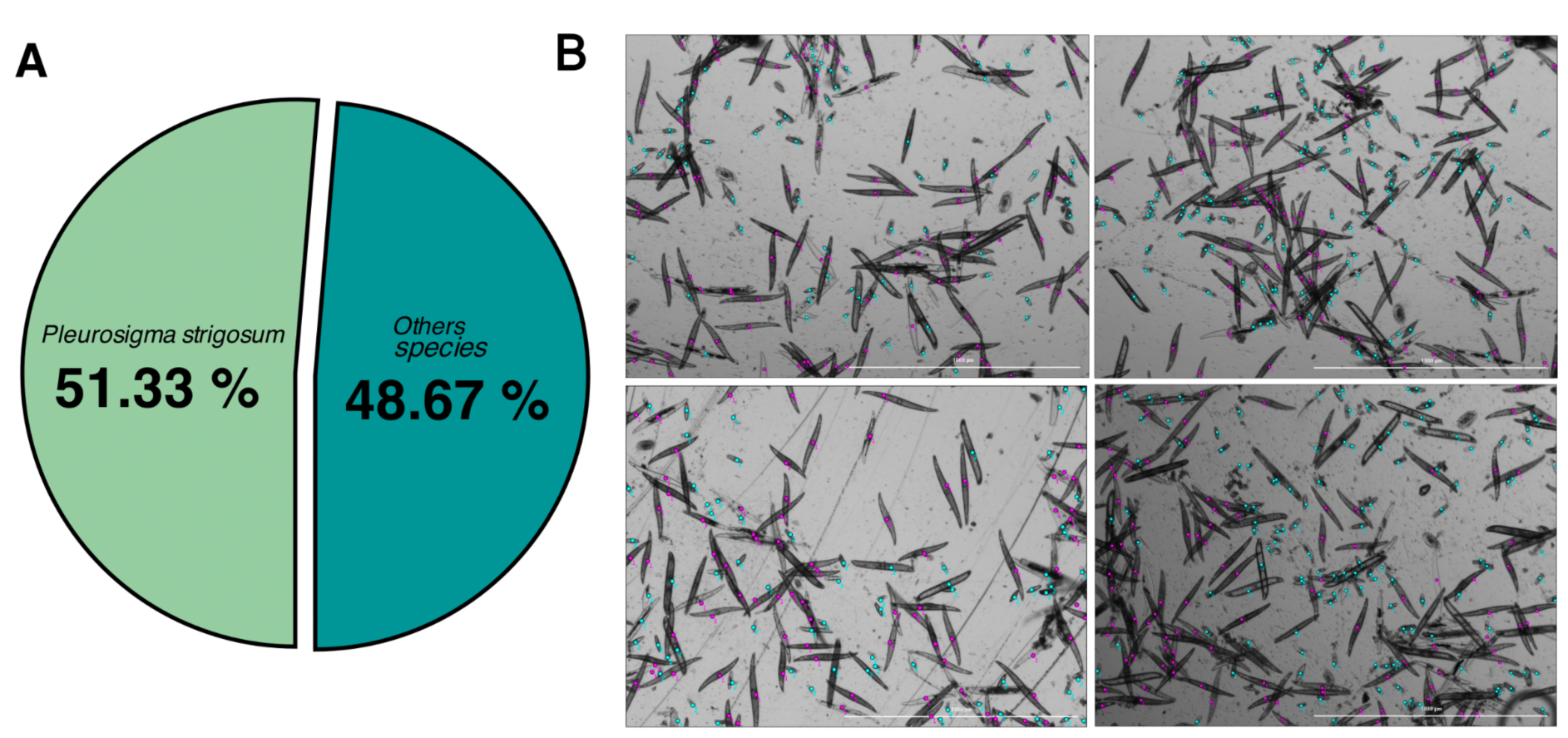
Diversity of the diatom assemblage. **A.** Average pie chart of diatom diversity (based on n = 10 images and, 1765 identified diatoms); **B.** Representative images used to assess diatom diversity.

**Figure s3.**
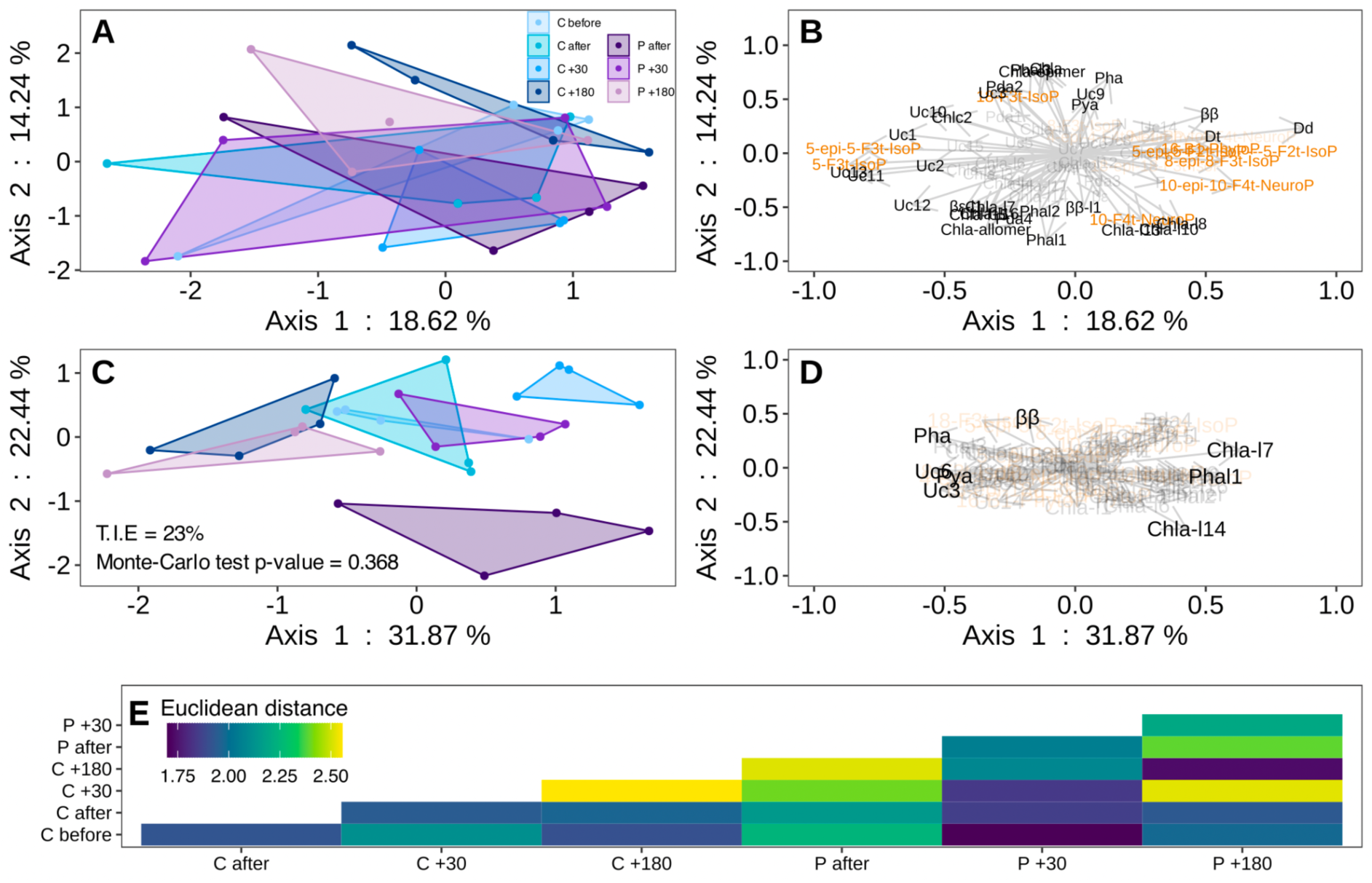
Lipophilic pigment and isoprostanoid responses to 2-min cold atmospheric plasma. **A.** Projection of samples in the factorial space derived from a Multiple Factor Analysis (MFA) based on their lipophilic pigment and isosprostanoid composition (%) across treatments; **B.** Projection of variable vectors in the MFA factorial space, showing only variables with cos^2^ > 0.2 for improved readability. Vectors directions and lengths indicate their correlations and relative contributions to the structuring of the factorial space; **C.** Between-class analysis performed on the MFA object (BC-MFA), constrained by the explanatory treatment variable. The total inertia explained (T.I.E.) reflects the proportion of variance associated with the partition imposed by the explanatory variable; **D.** Projection of variable vectors in the BC-MFA factorial space, showing only variables with cos^2^ > 0.2 for improved readability. Vectors directions and lengths indicate their correlations and relative contributions to the structuring of the factorial space; **E.** Heatmap of Euclidean distances between group centroids performed across all MFA dimensions. 2-min cold atmospheric plasma treatments (n = 4): Control before, C before; Control after, C after; Control +30 min after treatment cessation, C +30; Control +180 min after treatment cessation, C +180; Cold atmospheric plasma treated-samples after, P after; Cold atmospheric plasma treated-samples +30 min after treatment cessation, P +30; Cold atmospheric plasma treated-samples +180 min after treatment cessation, P +180. The abbreviations for pigment names are defined in Supplementary Table 2.

**Figure s4.**
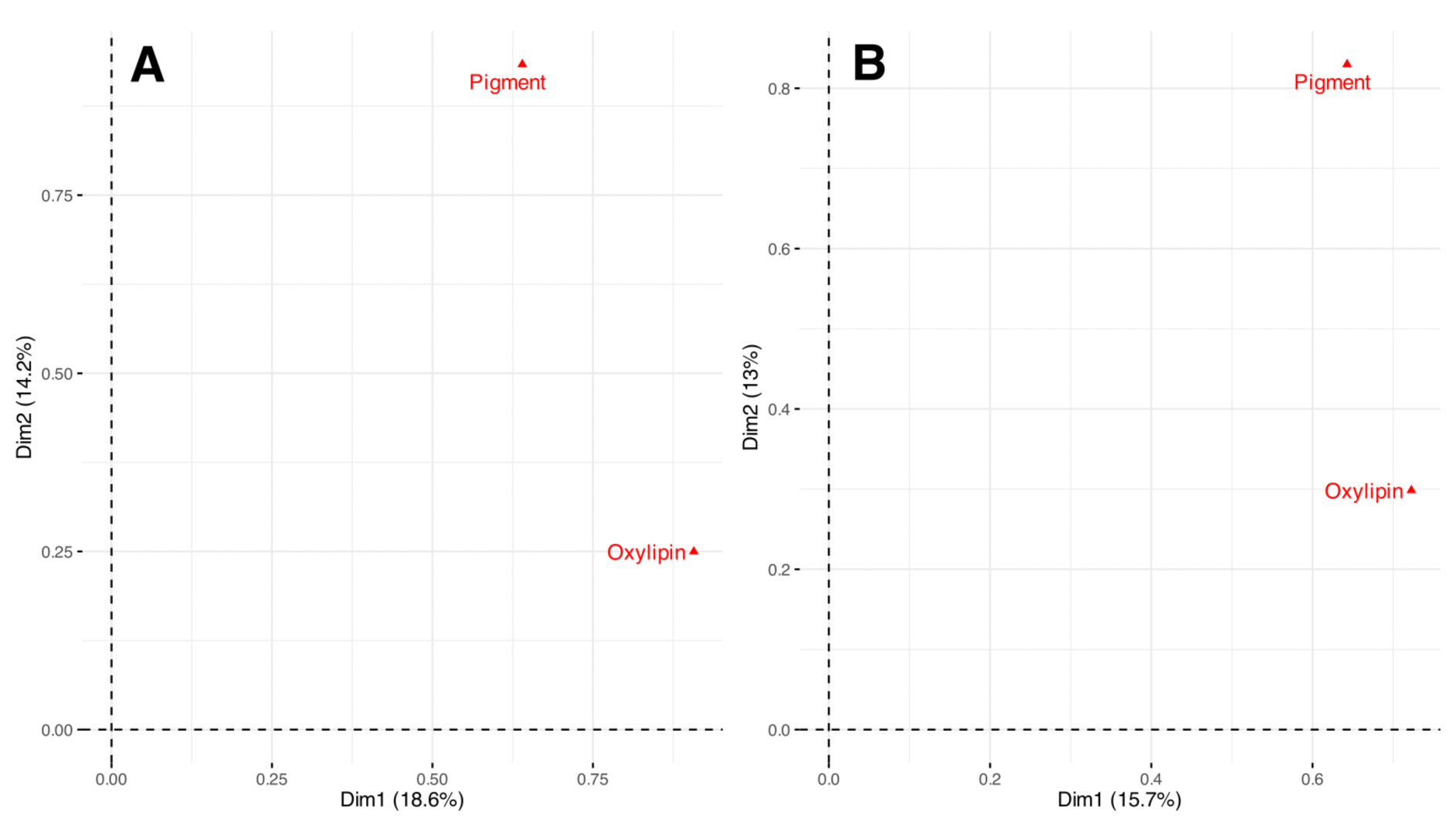
Pigment and isoprostanoid group representation in the multivariate MFA space. **A.** Pigment and isoprostanoid group representation following 2-min cold atmospheric plasma treatment; **B.**Pigment and isoprostanoid group representation following 5-min cold atmospheric plasma treatment.

**Figure s5.**
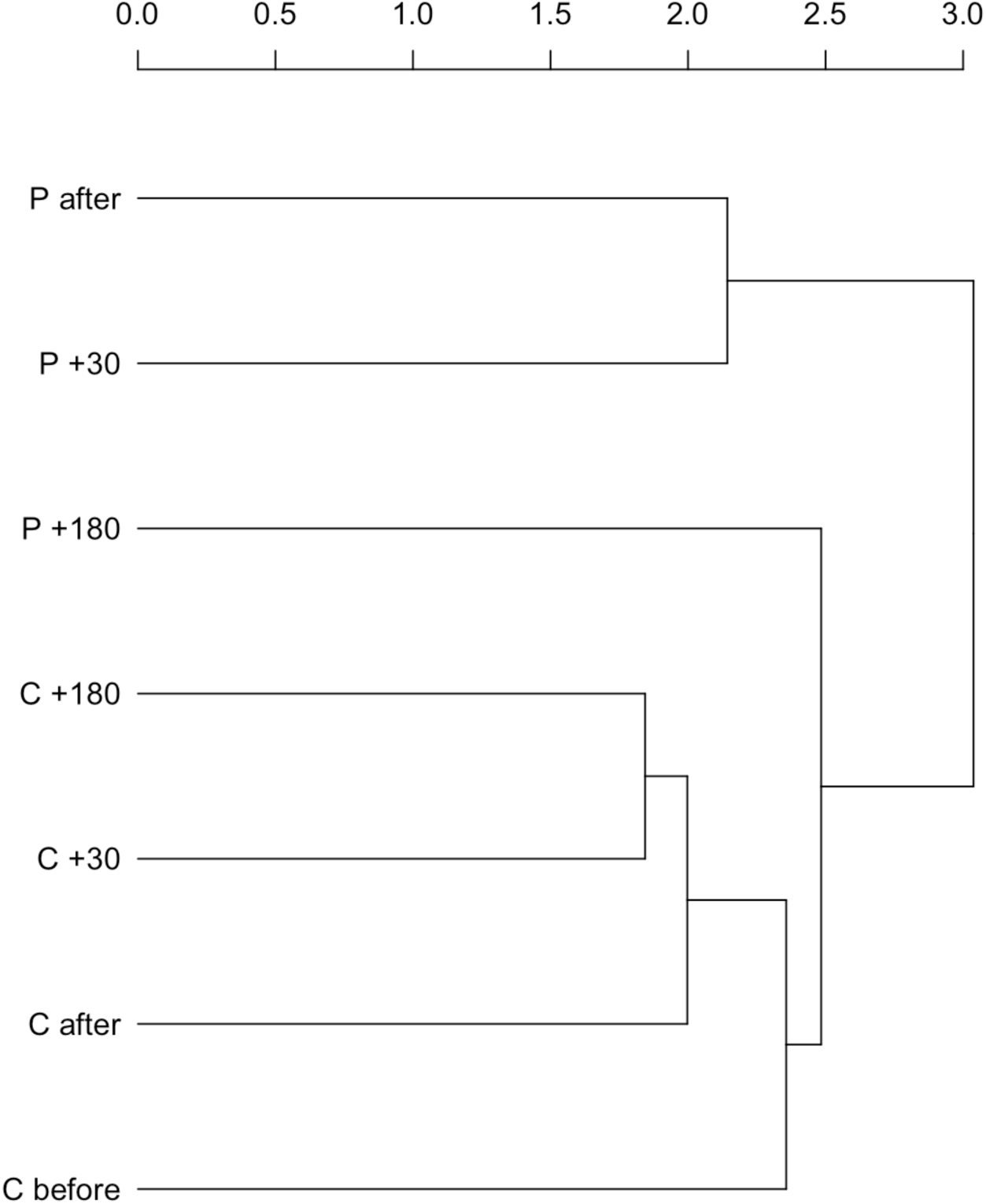
Hierarchical clustering of treatments based on distances between MFA group centroids. Hierarchical clustering of experimental treatments based on Euclidean distances between group centroids across all dimensions of the MFA factorial space, integrating pigment and isoprostanoid profiles.

**Figure s6.**
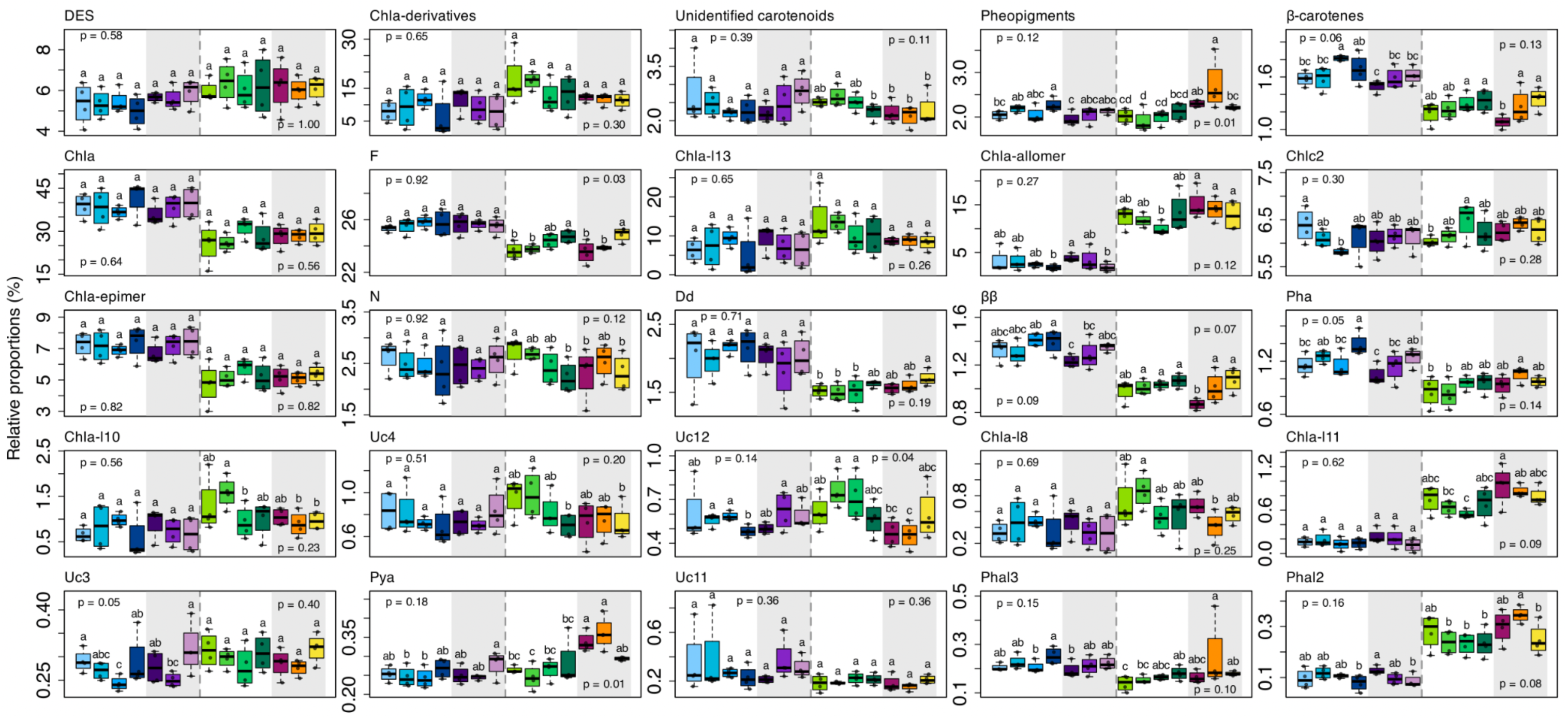
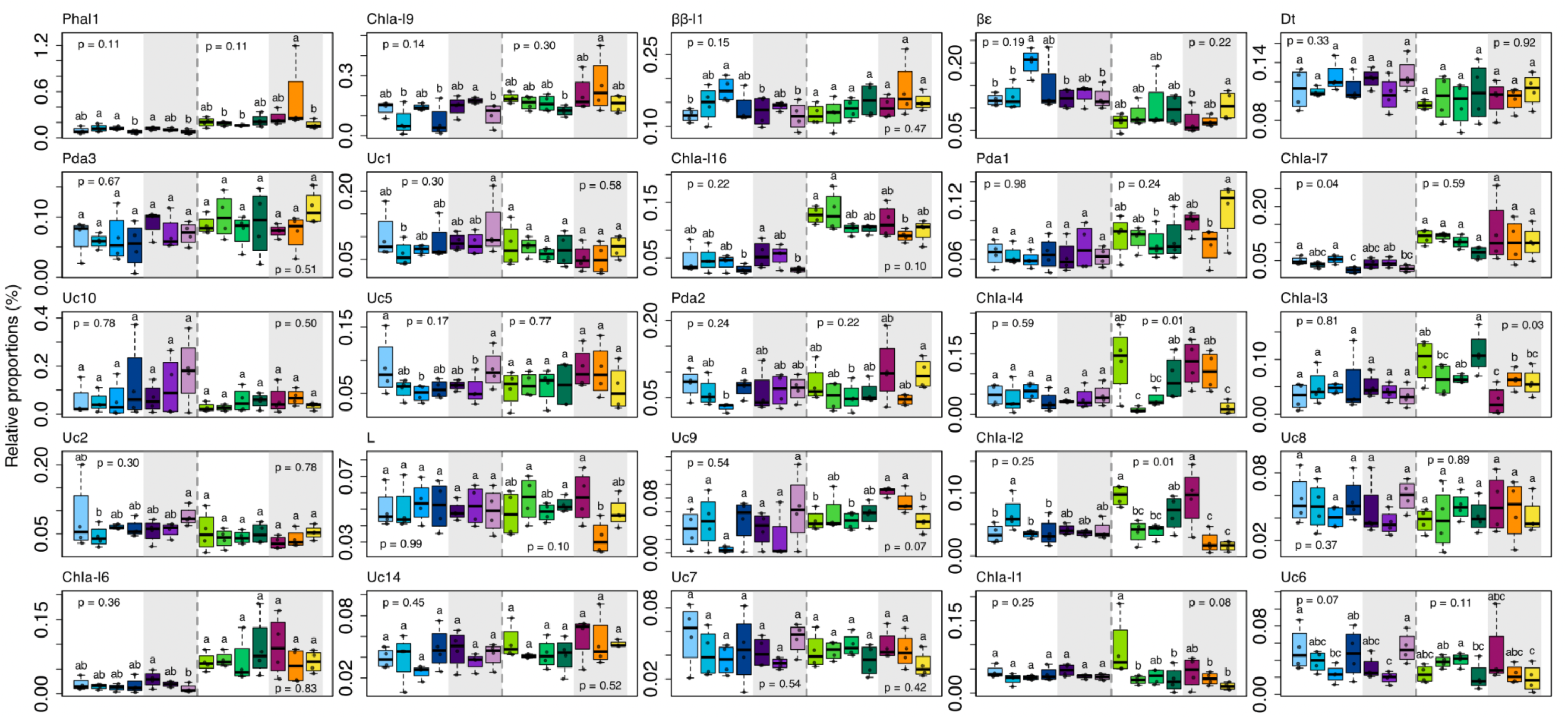
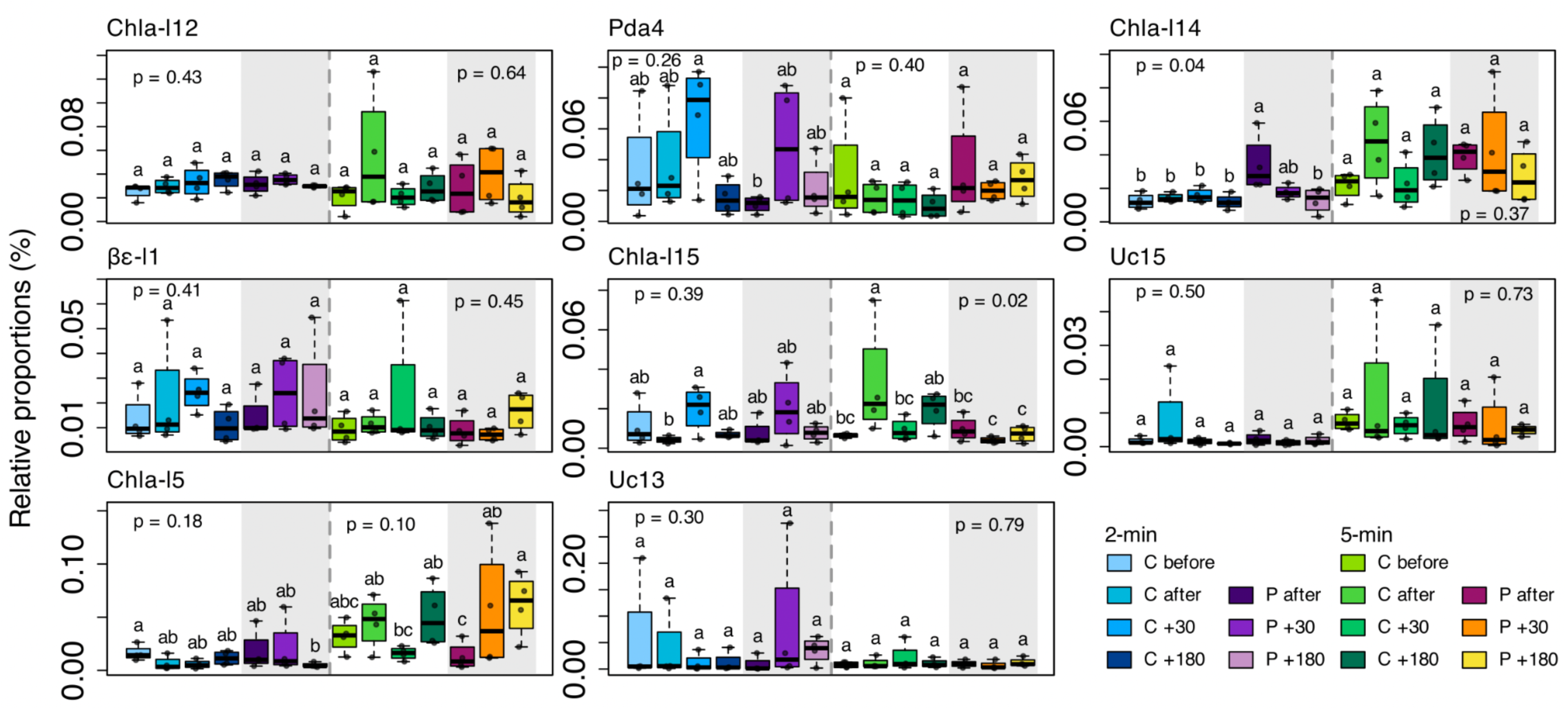
Boxplot of individual pigment changes under cold atmospheric plasma treatments. Boxplots showing the relative proportions (%, calculated from pigment concentrations) of individual lipophilic pigments under 2-min (left panel) and 5-min (right panel) cold atmospheric plasma treatments. Pigment boxplots are ordered by decreasing relative proportion across samples, except for the first five boxplot groups, which represent, respectively: the diadinoxanthin de-epoxidation state (diatoxanthin/diadinoxanthin + diatoxanthin × 100), DES; the chlorophyll *a*-derivatives group (Chla-derivatives), including the sum of all chlorophyll *a*-like forms; the unidentified carotenoids group, including the sum of all unidentified carotenoid pigments; the pheopigment group, including the sum of pheophytin *a*, pheophorbide *a*, and pyropheophytin *a*; and the β-carotenes group, including the sum of β-β-carotene and β-ε-carotene forms. 2-min and 5-min cold atmospheric plasma treatments: Control before, C before; Control after, C after; Control +30 min after treatment cessation, C +30; Control +180 min after treatment cessation, C +180; Cold atmospheric plasma treated-samples after, P after; Cold atmospheric plasma treated-samples +30 min after treatment cessation, P +30; Cold atmospheric plasma treated-samples +180 min after treatment cessation, P +180. In each 2-min and 5-min cold atmospheric plasma treatments, groups were statistically tested using the Van der Waerden test (n = 4). The abbreviations for pigment names are defined in Supplementary Table 2.

**Figure s7.**
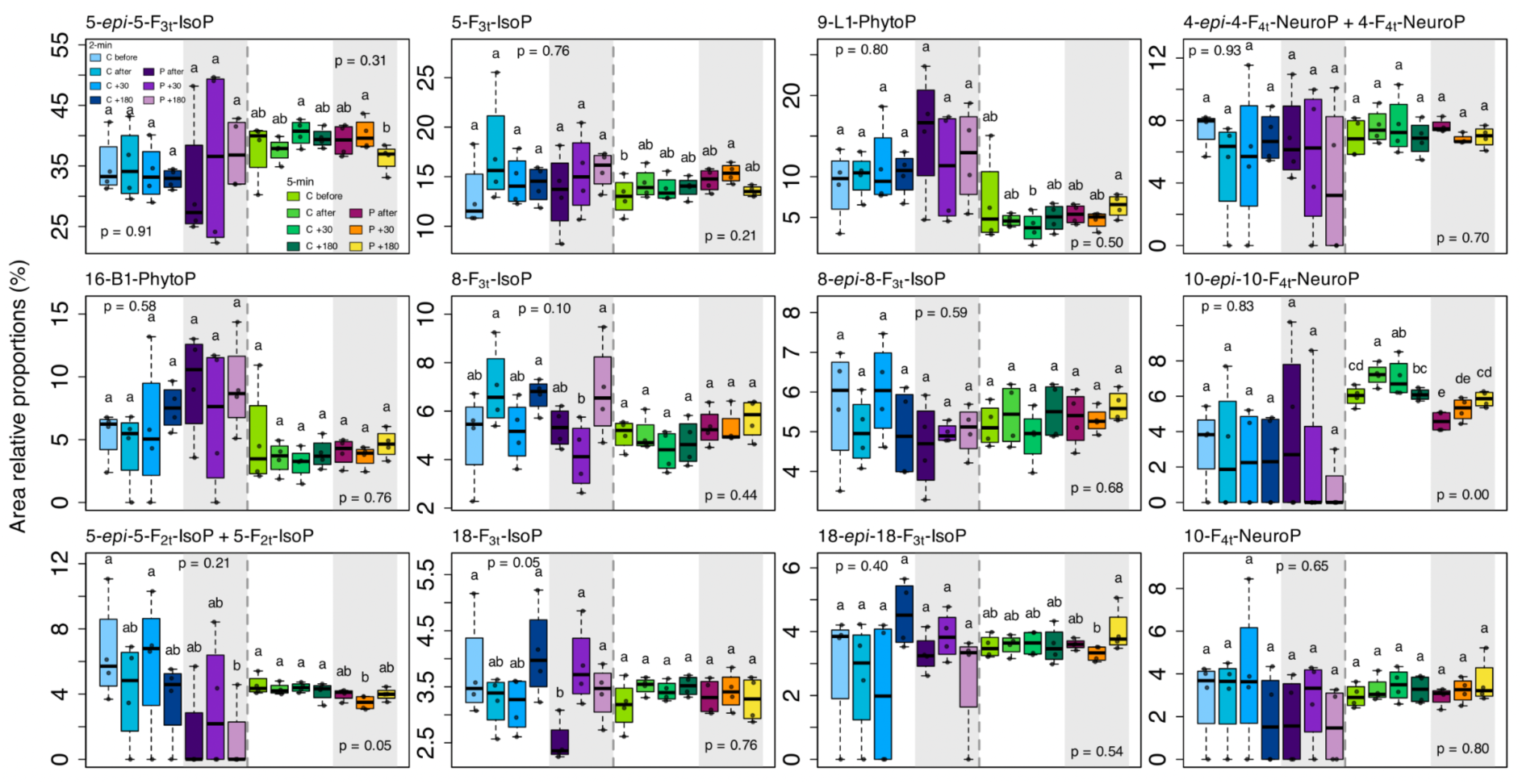
Boxplot of individual isoprostanoid changes under cold atmospheric plasma treatments. Boxplots showing the relative proportions (%, calculated from raw chromatogram area) of individual isoprostanoids under 2-min (left panel) and 5-min (right panel) cold atmospheric plasma treatments. Isoprostanoid boxplots are ordered by decreasing relative proportion across samples. 2-min and 5-min cold atmospheric plasma treatments: Control before, C before; Control after, C after; Control +30 min after treatment cessation, C +30; Control +180 min after treatment cessation, C +180; Cold atmospheric plasma treated-samples after, P after; Cold atmospheric plasma treated-samples +30 min after treatment cessation, P +30; Cold atmospheric plasma treated-samples +180 min after treatment cessation, P +180. In each 2-min and 5-min cold atmospheric plasma treatments, groups were statistically tested using the Van der Waerden test (n = 4).

**Figure s8.**
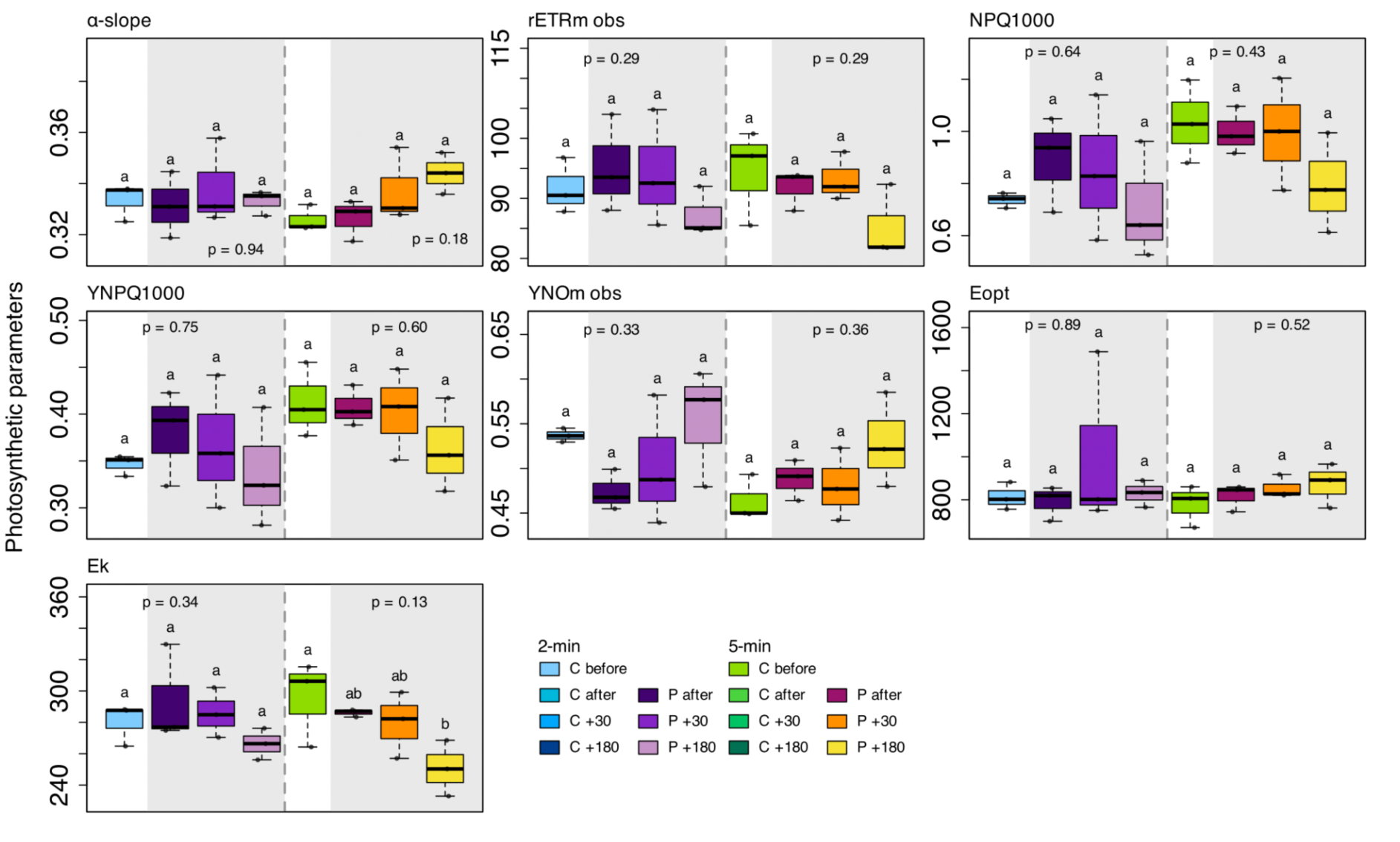
Boxplot of photosynthetic parameter changes under cold atmospheric plasma treatments. Boxplots showing photosynthetic parameter changes under 2-min (left panel) and 5-min (right panel) cold atmospheric plasma treatments. 2-min and 5-min cold atmospheric plasma treatments: Control before, C before; Cold atmospheric plasma treated-samples after, P after; Cold atmospheric plasma treated-samples +30 min after treatment cessation, P +30; Cold atmospheric plasma treated-samples +180 min after treatment cessation, P +180. In each 2-min and 5-min cold atmospheric plasma treatments, groups were statistically tested using the Van der Waerden test (n = 3).

---

**Table s1.**
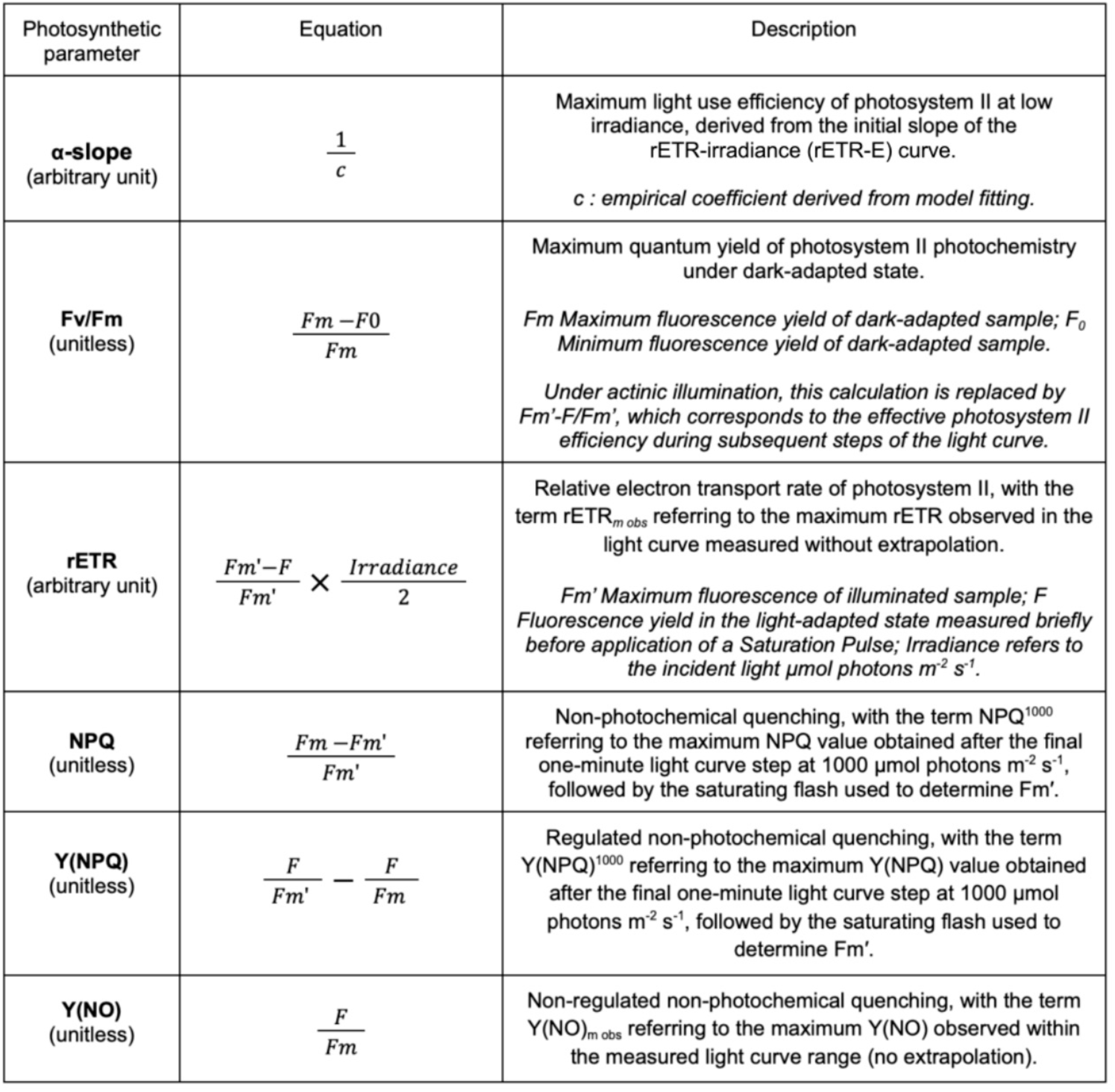
Photosynthetic parameters definitions. Equations adapted from Consalvey (2005), and Christof Klughammer and Ulrich Schreiber (2008) used to calculate photosynthetic parameters.

| Photosynthetic parameter | Equation | Description |
| --- | --- | --- |
| **α-slope**<br>(arbitrary unit) | $\frac{1}{c}$ | Maximum light use efficiency of photosystem II at low irradiance, derived from the initial slope of the rETR-irradiance (rETR-E) curve.<br>*c : empirical coefficient derived from model fitting.* |
| **Fv/Fm**<br>(unitless) | $\frac{F_m - F_0}{F_m}$ | Maximum quantum yield of photosystem II photochemistry under dark-adapted state.<br>*F<sub>m</sub> Maximum fluorescence yield of dark-adapted sample; F<sub>0</sub> Minimum fluorescence yield of dark-adapted sample.*<br>*Under actinic illumination, this calculation is replaced by F<sub>m</sub>'-F/F<sub>m</sub>', which corresponds to the effective photosystem II efficiency during subsequent steps of the light curve.* |
| **rETR**<br>(arbitrary unit) | $\frac{F_m' - F}{F_m'} \times \frac{\text{Irradiance}}{2}$ | Relative electron transport rate of photosystem II, with the term rETR<sub>m obs</sub> referring to the maximum rETR observed in the light curve measured without extrapolation.<br>*F<sub>m</sub>' Maximum fluorescence of illuminated sample; F Fluorescence yield in the light-adapted state measured briefly before application of a Saturation Pulse; Irradiance refers to the incident light $\mu\text{mol photons m}^{-2} \text{s}^{-1}$.* |
| **NPQ**<br>(unitless) | $\frac{F_m - F_m'}{F_m'}$ | Non-photochemical quenching, with the term NPQ<sup>1000</sup> referring to the maximum NPQ value obtained after the final one-minute light curve step at 1000 $\mu\text{mol photons m}^{-2} \text{s}^{-1}$, followed by the saturating flash used to determine F<sub>m</sub>'. |
| **Y(NPQ)**<br>(unitless) | $\frac{F}{F_m'} - \frac{F}{F_m}$ | Regulated non-photochemical quenching, with the term Y(NPQ)<sup>1000</sup> referring to the maximum Y(NPQ) value obtained after the final one-minute light curve step at 1000 $\mu\text{mol photons m}^{-2} \text{s}^{-1}$, followed by the saturating flash used to determine F<sub>m</sub>'. |
| **Y(NO)**<br>(unitless) | $\frac{F}{F_m}$ | Non-regulated non-photochemical quenching, with the term Y(NO)<sub>m obs</sub> referring to the maximum Y(NO) observed within the measured light curve range (no extrapolation). |

**Table s2.**
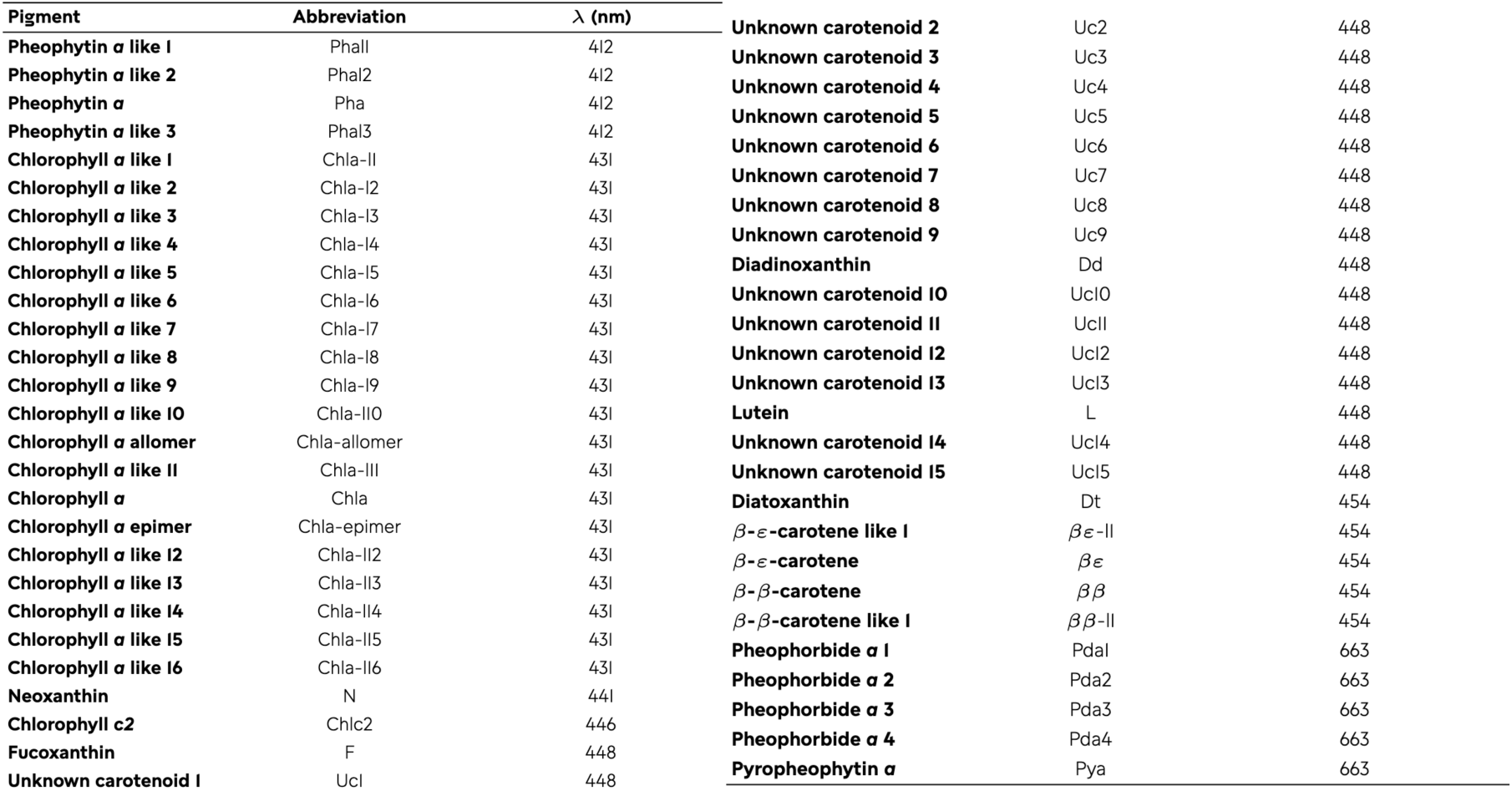
Pigment diversity in microalgal suspension samples.

**Table s3.** Table of raw photosynthetic parameters. Values (mean ± SD) before and after 2-min and 5-min cold atmospheric plasma treatment:: Control before, C before; Control after, C after; Control +30 min after treatment cessation, C +30; Control +180 min after treatment cessation, C +180; Cold atmospheric plasma treated-samples after, P after; Cold atmospheric plasma treated-samples +30 min after treatment cessation, P +30; Cold atmospheric plasma treated-samples +180 min after treatment cessation, P +180. ^(2)^ correspond to 2-min samples experiment and ^(5)^ correspond to 5-min samples experiment. α-slope (arbitrary unit) corresponds to the maximum light use efficiency of photosystem II under low irradiances. Fv/Fm (unitless) corresponds to the maximum quantum yield of photosystem II photochemistry under dark-adapted state. rETR_m obs_ (arbitrary unit) and Y(NO)_m obs_ (unitless) refer to the maximum values observed within the measured light curve range (without extrapolation). NPQ^1000^ (unitless) and Y(NPQ)^1000^ (unitless) correspond to the maximum values measured after the final one-minute light curve step at 1000 μmol photons m^-2^ s^-1^ (without extrapolation). Eopt refers to the optimal light parameter for autotrophic cells (μmol photons m^-2^ s^-1^). Ek refers to the light saturation coefficient where the cell begins to activate photoprotective mechanisms (μmol photons m^-2^ s^-1^). All fluorescence measurements were performed on dark-adapted samples. For each photophysiological parameter, mean ± SD values were calculated from triplicate (n = 3).

| Treatment | α-slope | Fv/Fm | $rETR_{m\ obs}$ | $NPQ^{1000}$ | $Y(NPQ)^{1000}$ | $Y(NO)_{m\ obs}$ | $E_{opt}$ | $E_k$ |
| --- | --- | --- | --- | --- | --- | --- | --- | --- |
| (2)C before | 0.33 (0.01) | 0.65 (0.00) | 91.69 (4.63) | 0.74 (0.03) | 0.35 (0.01) | 0.54 (0.01) | 813.20 (64.10) | 280.20 (13.37) |
| (2)P after | 0.33 (0.01) | 0.64 (0.02) | 95.17 (8.13) | 0.89 (0.18) | 0.38 (0.05) | 0.47 (0.02) | 791.23 (81.02) | 293.85 (31.12) |
| (2)P +30 | 0.34 (0.02) | 0.65 (0.01) | 94.30 (9.73) | 0.85 (0.28) | 0.37 (0.07) | 0.50 (0.07) | 1013.26 (411.95) | 285.82 (15.94) |
| (2)P +180 | 0.33 (0.00) | 0.66 (0.01) | 87.28 (4.09) | 0.71 (0.23) | 0.34 (0.06) | 0.55 (0.07) | 829.31 (62.59) | 266.22 (10.07) |
| (5)C before | 0.33 (0.01) | 0.65 (0.01) | 94.44 (7.98) | 1.03 (0.16) | 0.41 (0.04) | 0.46 (0.03) | 778.82 (97.77) | 295.29 (27.27) |
| (5)P after | 0.33 (0.01) | 0.62 (0.01) | 91.77 (3.35) | 1.00 (0.09) | 0.41 (0.02) | 0.49 (0.02) | 816.26 (63.19) | 286.12 (2.42) |
| (5)P +30 | 0.34 (0.01) | 0.66 (0.02) | 93.23 (4.05) | 0.99 (0.22) | 0.40 (0.05) | 0.48 (0.04) | 855.11 (54.09) | 279.47 (21.24) |
| (5)P +180 | 0.34 (0.01) | 0.66 (0.00) | 85.33 (6.08) | 0.79 (0.19) | 0.36 (0.05) | 0.53 (0.05) | 872.72 (103.32) | 250.58 (17.81) |

